# A Robust Machine Learning Framework for Keloid Biomarker Discovery Beyond Differential Expression

**DOI:** 10.64898/2026.06.24.734231

**Authors:** Ali Daher, Fareeha Afzal, Raluca Eftimie

## Abstract

Keloids are fibroproliferative skin disorders arising following dermal injury that extend beyond the original wound margins. Their pathogenesis remains poorly understood, and current treatments are associated with high recurrence rates. Identifying transcriptomic biomarkers that distinguish keloids from other skin and scar phenotypes may provide insight into disease mechanisms and facilitate the development of targeted therapeutic approaches. However, previous transcriptomic studies have often been limited by small sample sizes, pairwise comparisons between tissue classes, heterogeneous data-integration strategies, and a reliance on conventional differential gene expression (DGE) analysis. Here, we employed a multi-stage machine learning (ML) workflow for robust keloid biomarker discovery using transcriptomic datasets derived from both bulk RNA sequencing and single-cell RNA sequencing (scRNA-seq). We assembled and harmonized, to the best of our knowledge, the largest curated cross-study keloid transcriptomic cohort currently available, comprising 81 samples from 13 independent studies spanning four clinically relevant tissue classes: normal skin, normotrophic scar, hypertrophic scar, and keloid scar. Through study-aware cross-validation, feature selection, partition-stability analysis, and bootstrap validation across multiple ML classifiers, we identified a panel of eight highly consistent biomarkers capable of distinguishing keloid from non-keloid samples. These biomarkers were associated with dysregulation of extracellular matrix homeostasis, fibrosis-resolution pathways, vascular remodelling, and metabolic reprogramming. Comparison with conventional DGE analysis demonstrated substantial agreement while also highlighting important differences between the two approaches. In particular, *FASN* was consistently identified by the ML workflow as an upregulated discriminatory biomarker despite exhibiting weak, non-significant differential expression in the DGE analysis. Cell-type-specific analysis further supported this finding, revealing significant *FASN* upregulation in fibroblast and vascular endothelial populations. These results demonstrate that ML and DGE capture complementary aspects of transcriptomic variation. This study provides a robust strategy for cross-study transcriptomic biomarker discovery and identifies candidate genes and pathways for future mechanistic and therapeutic investigation in keloids.

**Author Summary:** Keloids are abnormal scars that continue to grow beyond the original wound and can be difficult to treat because they frequently recur after therapy. Although many studies have investigated the biology of keloids, the molecular mechanisms that distinguish them from other scar types remain incompletely understood. Identifying biomarkers involved in keloid formation may help inform improved treatment strategies. Previous transcriptomic studies have often been limited by small sample sizes and inconsistent analytical approaches. In this study, we combined gene-expression data from multiple independent studies to create, to the best of our knowledge, the largest cross-study transcriptomic collection available for keloid analysis. We then applied several machine learning approaches to identify genes that consistently distinguished keloids from other skin and scar phenotypes. The identified biomarkers were associated with extracellular matrix remodeling, fibrosis, vascular function, and cellular metabolism. One gene involved in fatty-acid synthesis, *FASN*, was repeatedly identified by the machine learning analyses despite being overlooked by conventional gene-expression methods. Additional single-cell analyses confirmed elevated *FASN* expression in specific cell populations within keloid tissue. More broadly, this work provides a strategy for discovering robust biomarkers from heterogeneous biological datasets and identifies molecular targets for future studies of keloid disease.

## 2 Introduction

Keloids are a unique fibroproliferative skin disorder that arises from aberrant wound healing, in which the ecessive collagen deposition extends beyond the boundaries of the original injury. Due to several shared phenotypic and molecular characteristics with certain tumour classes, keloids are often described as benign fibroproliferative dermal tumours. [1, 2] Both conditions exhibit pathological features such as local invasion into surrounding tissue, sustained cellular proliferation, resistance to apoptosis, recurrence following treatment, and the capacity for vascularization. Unlike malignant tumours, however, keloids do not arise spontaneously and lack metastatic potential. [1, 3]

Of particular relevance to keloid scars are hypertrophic scars, which represent another form of pathological wound healing characterized by excessive extracellular matrix (ECM) deposition and fibrosis. In contrast to keloids, hypertrophic scars generally remain confined to the original wound margins and may undergo partial regression over time. [4, 5] The two scar types also differ in their temporal development, anatomical distribution, and epidemiological characteristics: whereas hypertrophic scars typically develop within weeks following injury and may gradually regress, keloids can emerge months after the initial wound, rarely regress spontaneously, and exhibit a predisposition for anatomical sites believed to be exposed to elevated mechanical tension, including the chest, shoulders, upper back, and earlobes. [6, 7] Keloids are also more frequently observed in individuals with darker skin pigmentation and often display familial predisposition. [4, 5]

Current treatment strategies for keloids include intralesional and topical therapies, surgical excision, radiation therapy, and laser-based interventions. [1, 8] Despite these approaches, recurrence remains common and treatment outcomes are often unpredictable. [8] A better understanding of the transcriptomic and proteomic profile of keloid scars can shed light on the mechanisms and biological pathways underlying their etiology and evolution, which may in turn inform the development or selection of more effective treatment approaches.

Previous studies have attempted to identify keloid biomarkers by comparing the transcriptomic profiles of keloid and non-keloid skin samples, predominantly through differential gene expression (DGE) analysis and, more recently, through machine learning (ML) approaches; see [7, 9–18], among others. However, several methodological challenges have limited, and continue to limit, the utility, reproducibility, and biological interpretability of transcriptomic biomarker discovery efforts in keloid pathology. One important limitation arises from the simultaneous inclusion of whole-tissue transcriptomic profiles and datasets generated from isolated cell populations, such as fibroblasts or immune cells, in the same comparative study. [16] Whereas isolated-cell datasets represent only a specific cellular compartment, whole-tissue samples capture the combined transcriptional contributions of all cell types present within the tissue. Consequently, differences identified by ML algorithms or DGE analyses may reflect the variation in cellular composition rather than disease-specific biology. Such discrepancies cannot be fully resolved through conventional batch-correction procedures because they originate from biological sampling differences rather than technical variation.

Additional heterogeneity arises from the integration of different transcriptomic technologies. For instance, previous meta-analyses have combined RNA sequencing datasets with microarray datasets, particularly Affymetrix microarrays. [16, 18] While both platforms quantify gene expression, they rely on fundamentally different measurement principles, units (hybridization intensities versus sequencing counts), statistical distributions, dynamic ranges, and noise characteristics. As a result, the corresponding differences extend beyond conventional batch effects and may introduce systematic biases that are difficult to eliminate through standard harmonization approaches. In contrast, bulk RNA-seq and pseudo-bulked scRNA-seq datasets share a common count-based sequencing framework, enabling more principled integration following appropriate normalization and batch correction procedures.

A further challenge may arise when animal (*e.g.,* murine) and human transcriptomic datasets are analysed within the same framework. [19] Fundamental biological differences exist between species, particularly with respect to wound-healing mechanisms. For example, murine wound closure is predominantly contraction-driven, whereas human wound healing relies more heavily on granulation tissue formation and re-epithelialization. [20] Consequently, pathways involved in scar formation and tissue remodelling may exhibit substantially different transcriptional signatures across species, limiting the biological interpretability and translational relevance of cross-species biomarker discovery efforts.

In addition, most transcriptomic studies focus on pairwise comparisons between only two tissue classes, such as keloid versus normal skin, [11] keloid versus normal scar, [14] or keloid versus hypertrophic scar. [9] While such comparisons can identify genes that distinguish the selected pair of conditions, they do not necessarily identify biomarkers uniquely associated with keloid pathology. For instance, genes identified through a keloid-versus-normal-skin comparison may reflect general wound-healing responses, ECM remodelling, or inflammatory processes that are shared across multiple scar phenotypes. Similarly, genes identified through comparisons between keloids and normal scars may capture transcriptomic signatures associated more broadly with fibrosis than with keloid disease specifically. Comparisons between keloids and hypertrophic scars are particularly informative because both conditions share many fibroproliferative and inflammatory features while exhibiting markedly different clinical behaviour. Such analyses may therefore help identify molecular mechanisms underlying features unique to keloids, including persistent growth beyond the original wound margins and resistance to treatment. However, studies restricted to these two scar types often involve very small sample sizes, thereby undermining their statistical power, and remain optimized for discrimination between a single pair of conditions. As a result, biomarkers derived from pairwise analyses may exhibit reduced specificity when evaluated across a broader spectrum of pathological and non-pathological skin states. The inclusion of multiple clinically relevant tissue classes within the same comparative study could, in principle, better enable the identification of biomarkers that more consistently distinguish keloids from skin and scar phenotypes.

Finally, the last few years have witnessed a rapid expansion in publicly available transcriptomic datasets, including numerous bulk RNA-seq and scRNA-seq studies of keloid and other scar pathologies. However, many of these recently published datasets have not been incorporated into previous integrative analyses. By incorporating the new datasets and leveraging both bulk RNA-seq and pseudo-bulked scRNA-seq data within a common count-based sequencing framework, it is possible to construct substantially larger transcriptomic cohorts than have previously been available. Such integration has the potential to increase statistical power and improve the robustness and generalizability of derived biological conclusions, which is particularly important given the historically small sample sizes of comparative transcriptomic studies investigating keloid-specific biomarkers, many of which contain fewer than ten samples in total.

To address the aforementioned limitations, the present study assembles, to the best of the authors’ knowledge, the largest curated transcriptomic cohorts of keloid and scar pathology currently available, comprising 81 samples collected from 13 independent studies and spanning four clinically relevant tissue classes: keloid scar, hypertrophic scar, normotrophic scar, and normal skin. By integrating bulk RNA-seq and pseudo-bulked scRNA-seq datasets, explicitly accounting for study-specific, platform-specific, and sequencing-chemistry effects, and employing multiple complementary ML paradigms for classification, we seek to identify transcriptomic biomarkers that are robust across independent studies, sequencing technologies, dataset partitioning strategies, and ML classifier architectures.

## 3 Results

### 3.1 Exploratory Analysis and Data Harmonization

The projection of the samples prior to harmonisation onto the three major PCs is shown in Figure 1. As shown in Figures 1(a) and 1(b), samples exhibited three distinct large-scale separations, which correspond to the first bulk RNA-seq experiment (GSE188592), the second bulk RNA-seq experiment (OEP00002674), and the remaining scRNA-seq-derived pseudo-bulk samples. Furthermore, within the scRNA-seq cohorts, another distinct but less pronounced variation emerged as a result of the sequencing chemistry/kit (Figure 1(c)), and to a lesser extent, as a result of the experimental study (Figure 1(b)). In contrast, no discernable separation or clustering pattern was observed based on scar/skin condition, with samples from different scar categories displaying substantial overlap and heterogeneous mixing in the PCA space in Figure 1(d).

**Figure 1:**
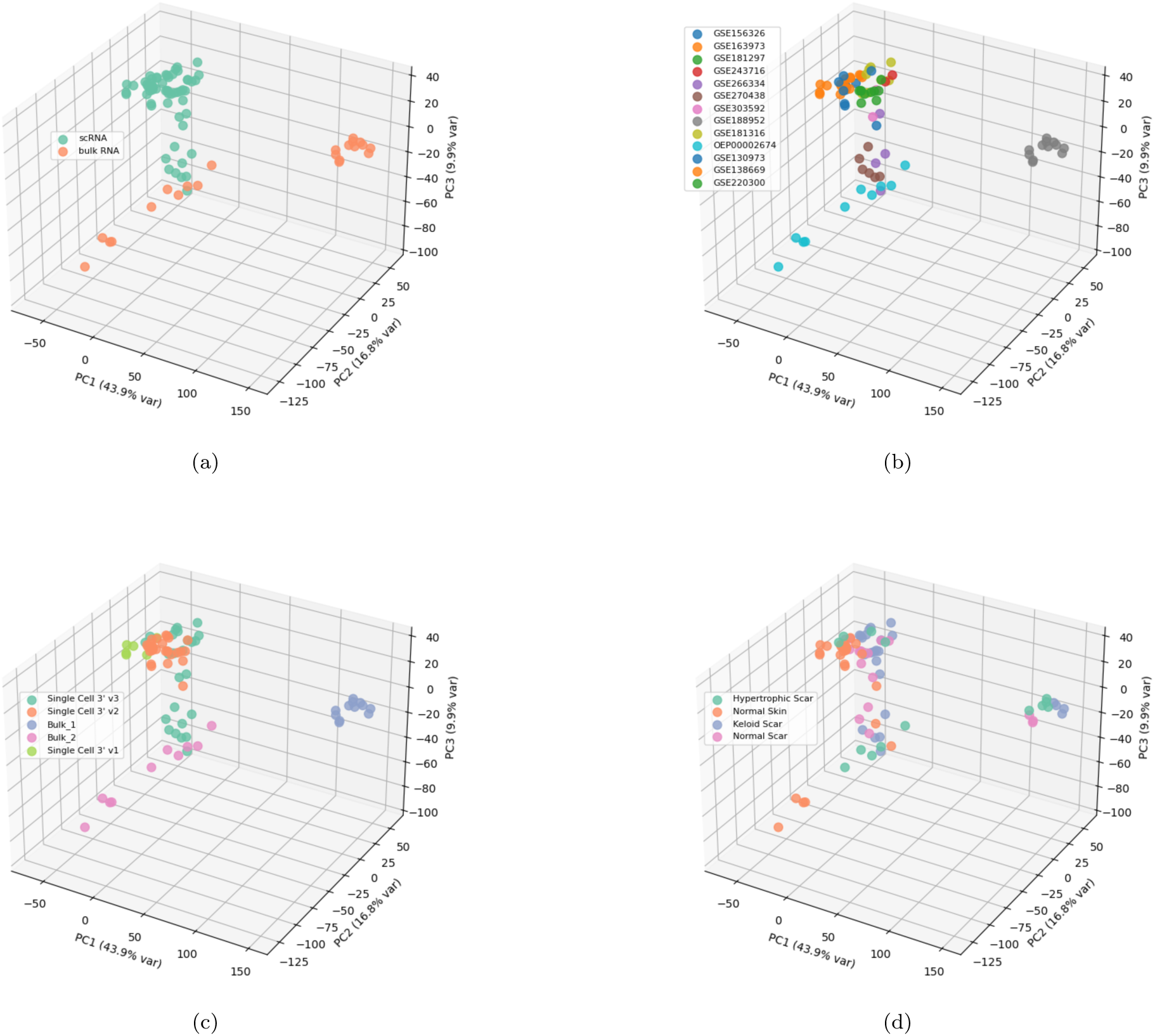
PCA Exploratory Analysis of the 81 Samples from Table : the log-normalized gene expression data were projected onto the first three PCs (of highest variance) and color-coded according to a)platform (scRNA vs bulk RNA), b)study (experiment), c)chemistry, and d)skin/Scar Category.

Figure 2(c) demonstrates that, following correction for technical covariates, the samples no longer exhibit strong clustering according to sequencing platform, study origin, or scRNA-seq chemistry (Figures 2(a), 2(b), and 2(c)). Instead, the corrected expression space begins to reveal biologically meaningful low-dimensional structure in the linearly transformed PCA space before the implementation of any ML classifiers, particularly in regard to the separation pattern between the normal skin and the rest of the scar samples.

**Figure 2:**
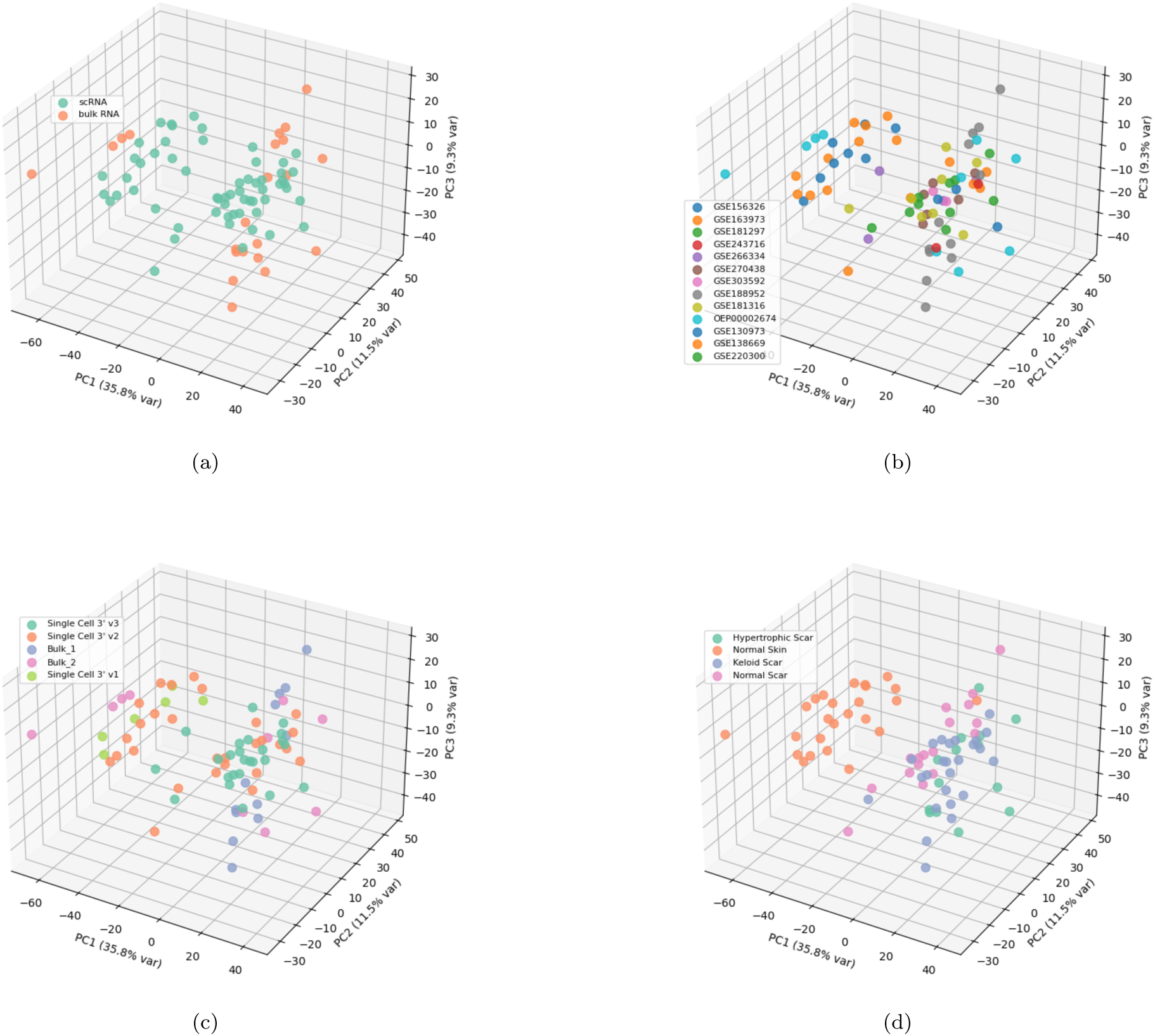
PCA Exploratory Analysis of the 81 Samples from Table 3: the log-normalized gene expression data were harmonized to subtract signal from technical variates and projected onto the first three PCs (of highest variance) and color-coded according to a)platform (scRNA vs bulk RNA), b)study (experiment), c)chemistry, and d)skin/Scar Category.

### 3.2 Classification Performance Across the Three Outer Folds

The mean and standard deviation of the accuracy and *F*_1_-scores across the outer cross-validation folds for each of the five ML algorithms are summarised in Table 1. The confusion matrices across the outer folds for the multiclass and binary classifiers are shown in Figures 3 and 4, respectively.

**Figure 3:**
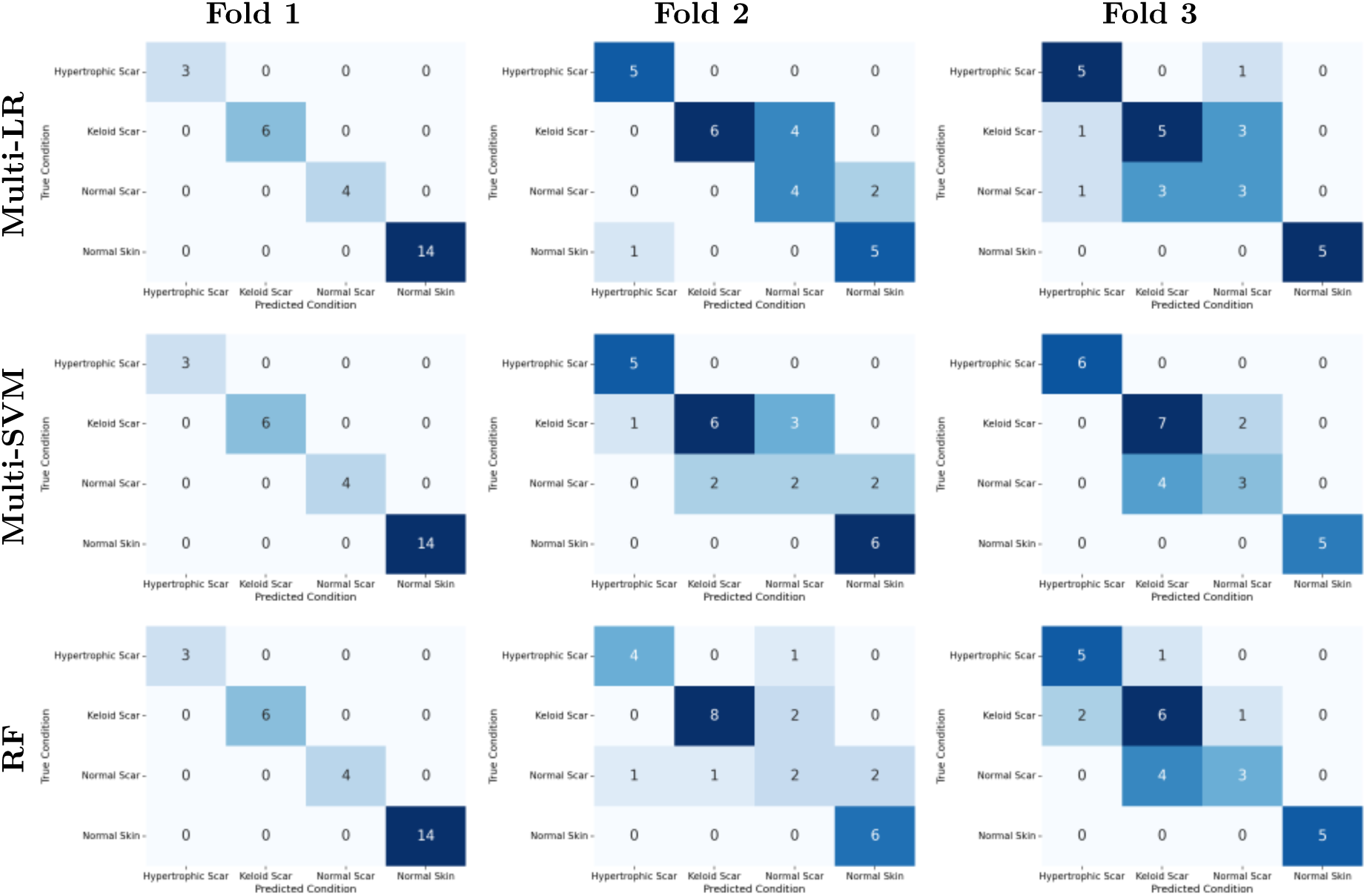
Multiclass confusion matrices across the three outer cross-validation folds.

**Figure 4:**
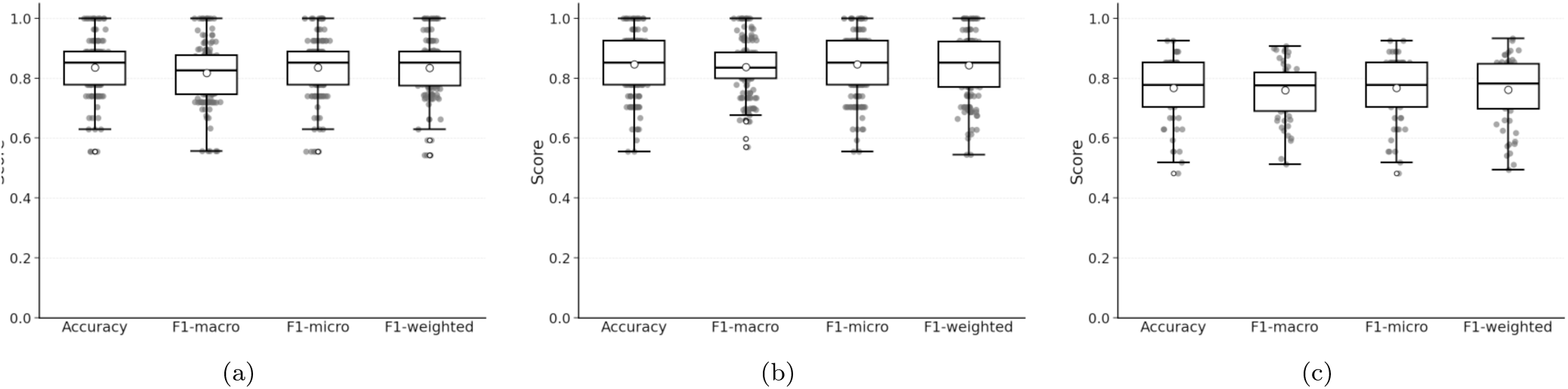
Binary confusion matrices across the three outer cross-validation folds.

**Table 1:** Classification performance across outer cross-validation folds. Values are reported as mean ± standard deviation.

| Binary classifiers |  |  | Multiclass classifiers |  |  |
| --- | --- | --- | --- | --- | --- |
| Algorithm | Accuracy | $F_1$ | Algorithm | Accuracy | Macro-$F_1$ |
| Bin-LR | $0.89 \pm 0.09$ | $0.86 \pm 0.11$ | Multi-LR | $0.81 \pm 0.14$ | $0.82 \pm 0.13$ |
| Bin-SVM | $0.88 \pm 0.09$ | $0.82 \pm 0.14$ | Multi-SVM | $0.83 \pm 0.13$ | $0.83 \pm 0.13$ |
| | | | RF | $0.81 \pm 0.13$ | $0.82 \pm 0.13$ |

### 3.3 Partition Stability Analysis Across Permissible Outer Folds

The resulting distributions of accuracy, macro-*F*_1_, micro-*F*_1_, and weighted-*F*_1_ scores for the multiclass classifiers are presented in Figure 5, while the corresponding distributions of accuracy, *F*_1_-score, precision, and recall for the binary classifiers are shown in Figure 6.

**Figure 5:**
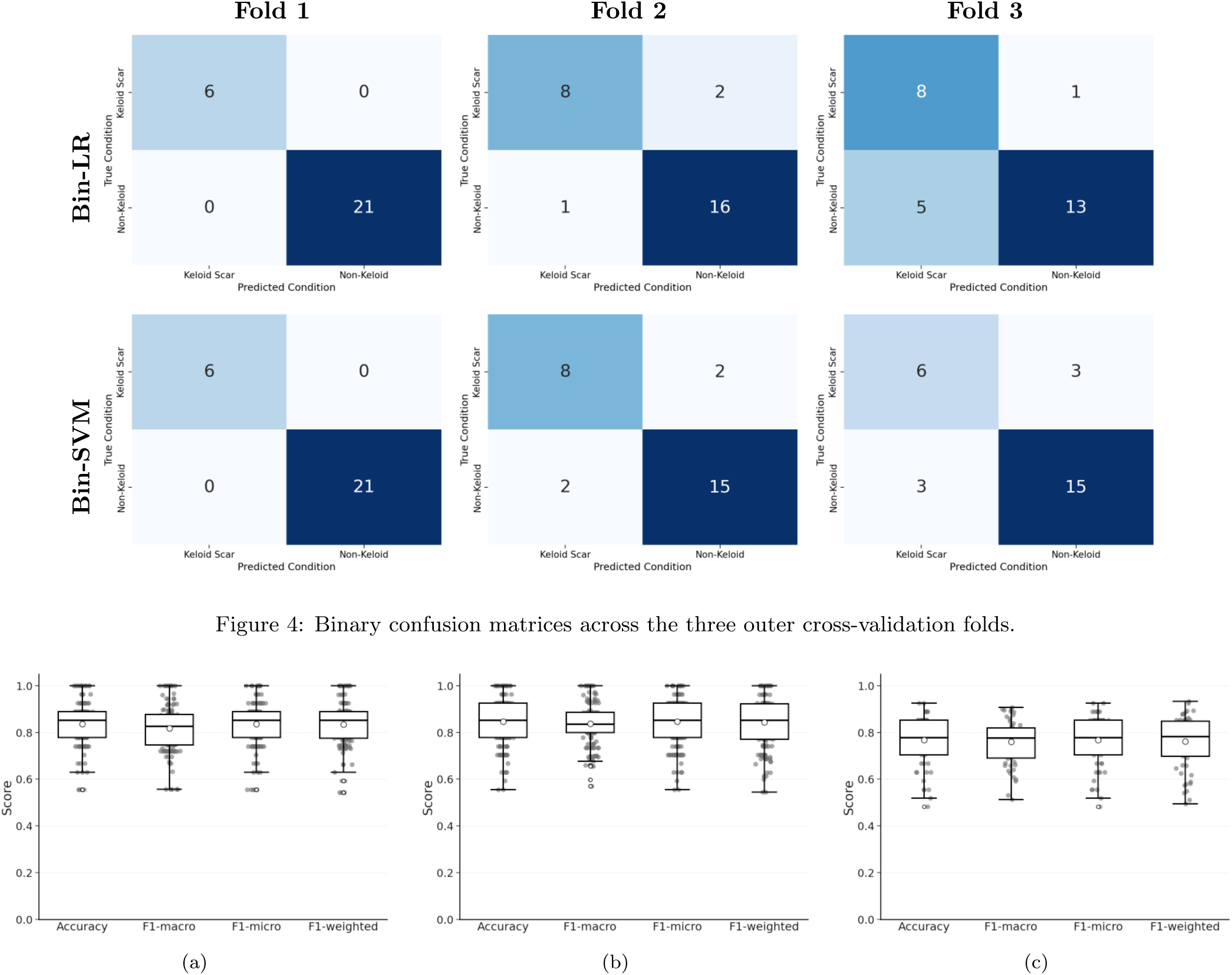
Distribution of multiclass classification performance metrics across the 56 eligible fold partitionings. (a) Multiclass logistic regression. (b) Multiclass linear support vector machine (SVM). (C) Random forest classifier. Boxplots summarise the distributions of accuracy, macro-*F*_1_, micro-*F*_1_, and weighted-*F*_1_ scores across study-aware partitionings.

**Figure 6:**
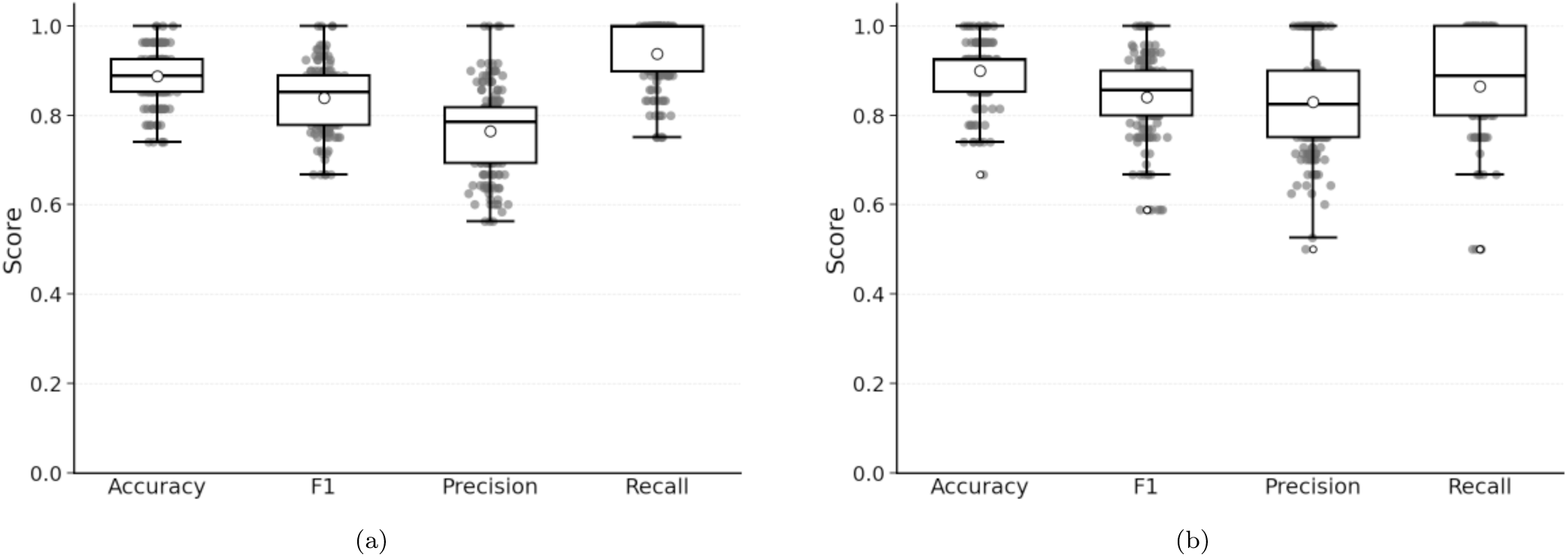
Distribution of binary classification performance metrics across the eligible 56 fold partitionings. (A) Binary logistic regression. (B) Binary linear support vector machine (SVM). Boxplots summarize the distributions of accuracy, *F*_1_ score, precision, and recall across study-aware partitionings.

### 3.4 Stable Biomarkers Identified Across Classifiers and Folds

Figure 7 summarises the resulting stable biomarker sets identified by each ML framework. Applying the criteria outlined in Section 5.7 yielded six consensus downregulated genes: *PCSK1N*, *CLEC3B*, *TNXB*, *AKR1C2*, *IL1RN*, and *SDC4* and eight consensus upregulated genes: *GATA2*, *KCNE4*, *FASN*, *BGN*, *CHN1*, *CDH11*, *ASPN*, and *CFI*.

**Figure 7:**
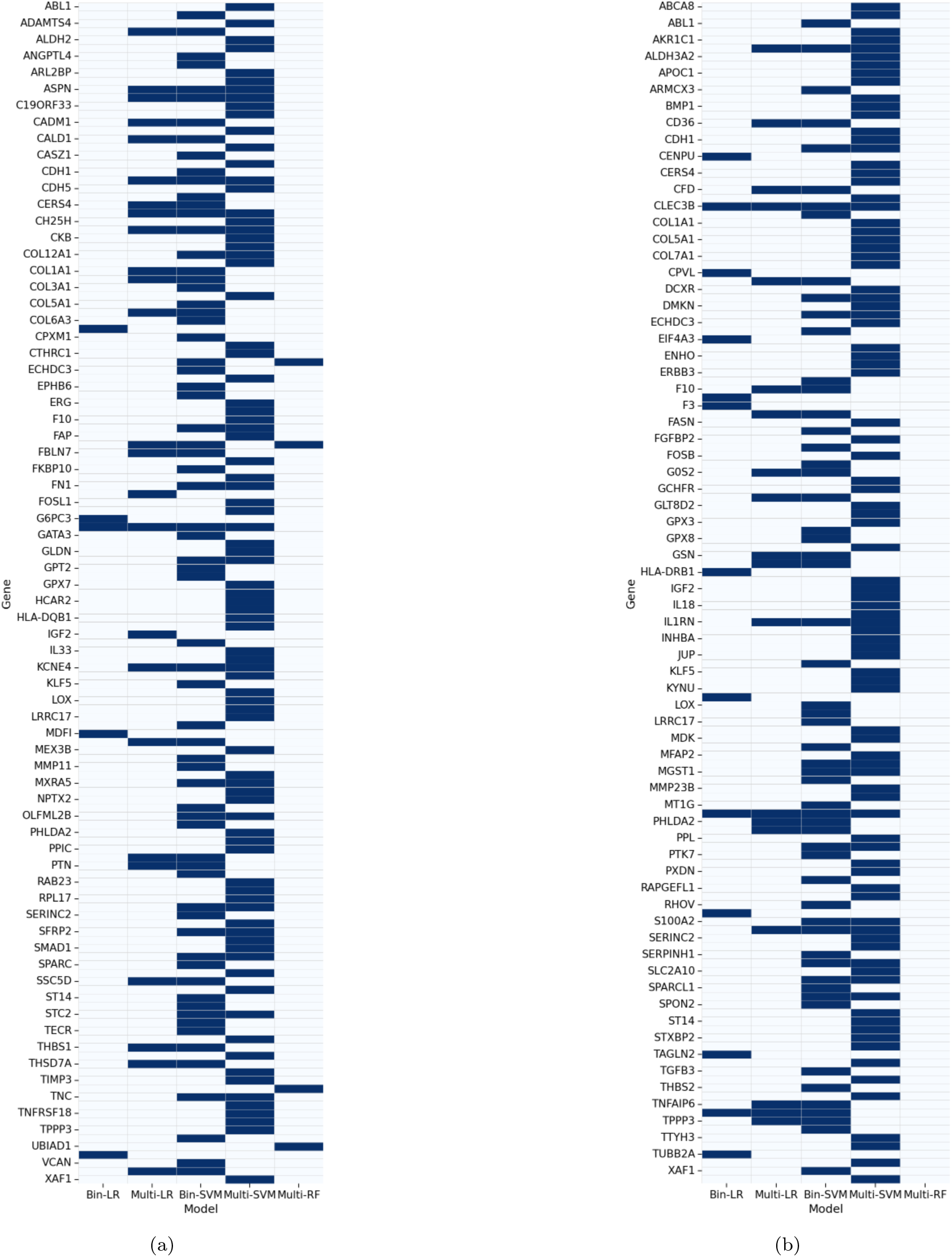
Stable keloid-associated transcriptomic biomarkers identified across the ML frameworks for (a) upregulated candidate biomarkers and (b) downregulated candidate biomarkers. Dark cells indicate that a gene was selected as stable for the corresponding ML classifier, meaning ≥ 70% representation across the eligible outer-fold partitionings.

### 3.5 Bootstrap Validation Analysis Using the Full Cohort of the Candidate Biomarkers

The 14 genes identified from Section 3.4 were used as features in a final step using a binary logistic regression model fitted on the full harmonized dataset. The final model achieved an accuracy of (0.94 ± 0.05) and an *F*_1_-score of (0.91 ± 0.07) across the CV inner folds and bootstrap simulations, indicating that the consensus biomarker panel retained strong discriminatory power across repeated resampling of the dataset. The coefficient distributions and corresponding confidence intervals for the 14 consensus biomarkers identified through the ML stability analysis are shown in Figure 8. Following bootstrap validation, of the 14 candidate biomarkers, five upregulated genes and three downregulated genes retained statistically significant regulatory directionality (*p <* 0.05).

**Figure 8:**
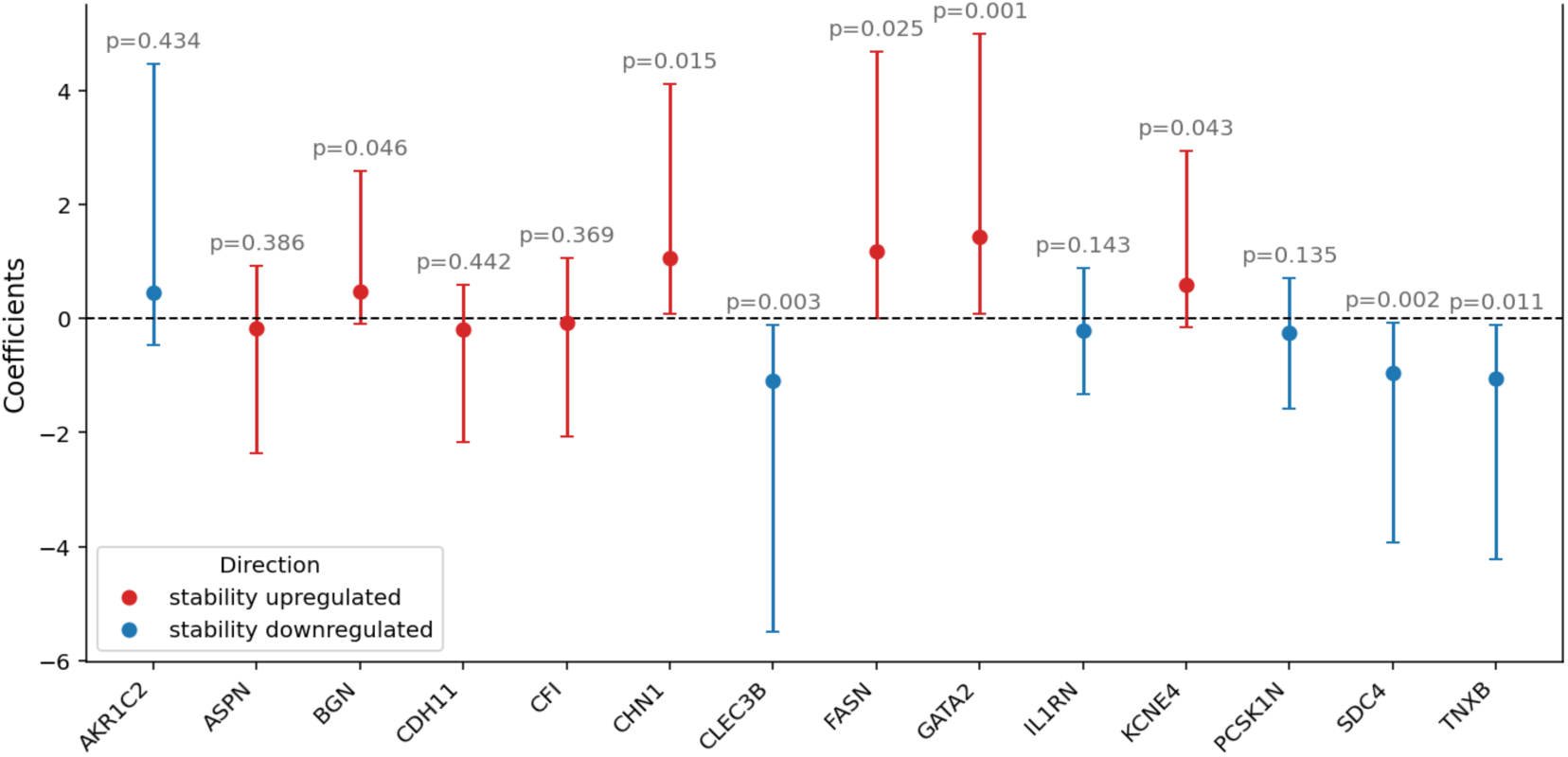
Binary logistic regression coefficient estimates, 95% percentile confidence intervals, and one-sided (p)-values for the 14 consensus biomarkers identified through the ML stability analysis when running 1000 bootstrap resample simulations on the full harmonized dataset.

### 3.6 Differential Gene Expression Analysis

The *log*_2_ fold changes and multiple-testing-adjusted (p)-values for the eight consensus biomarkers were extracted and are summarized in Table 2. Figure 9(a) presents a volcano plot demonstrating the *log*_2_ fold changes and adjusted-(p)-values of the genes used in the downstream IPA analysis. Figure 9(b) presents a bubble-volcano representation of selected pathways, in which bubble size reflects the number of overlapping molecules associated with each pathway, thereby illustrating the relationship between pathway significance, predicted activation state, and pathway coverage. Figure 9(c) summarises the highest-ranking upregulated canonical pathways according to IPA-predicted activation *z*-scores, while Figure 9(d) shows the most significantly enriched pathways ranked by adjusted *p* value. To provide an independent pathway-level validation of the IPA findings, Gene Set Enrichment Analysis (GSEA) [21] was also performed on the full ranked DESeq2 gene list, the results of which are provided in Figure S1.

**Figure 9:**
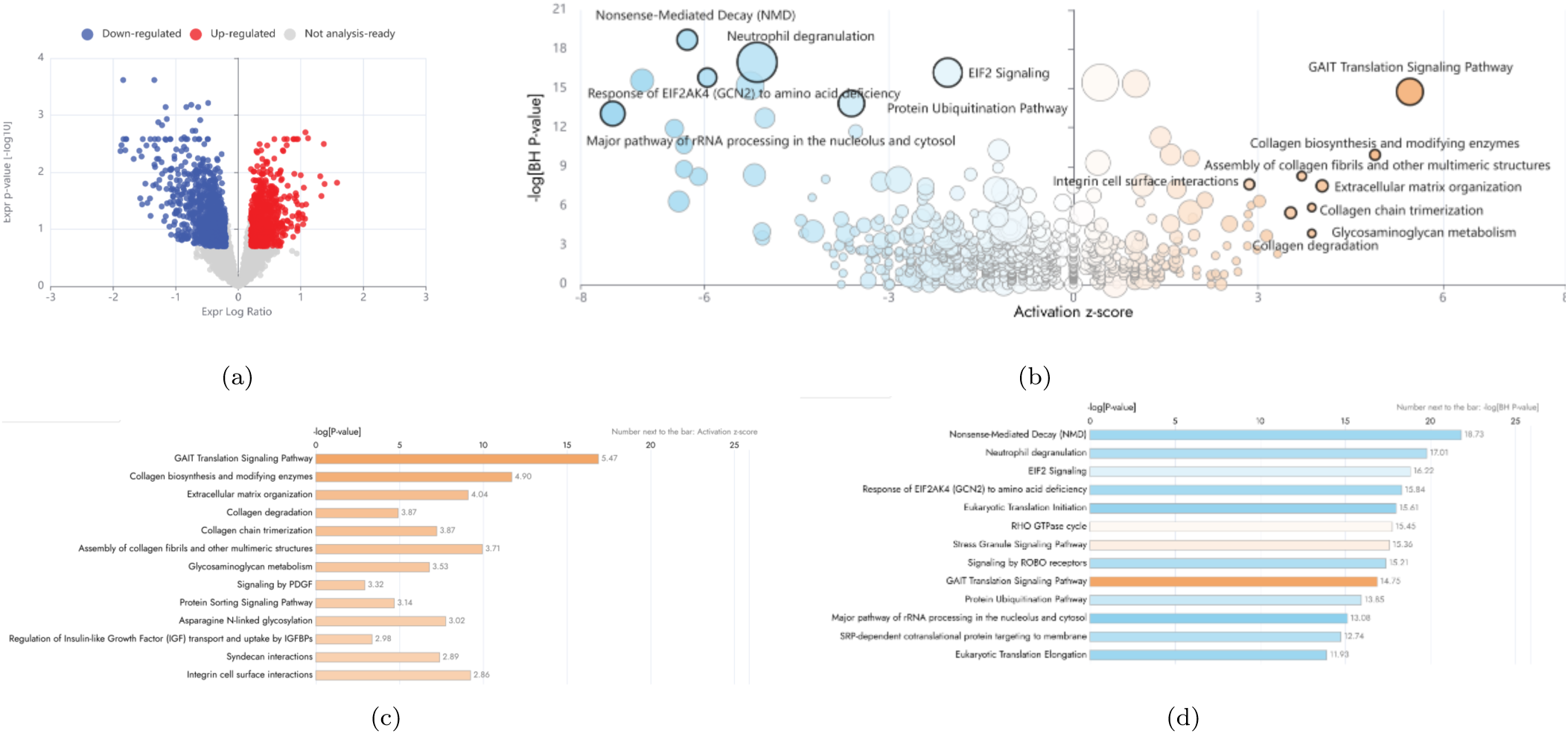
Differential gene expression and pathway enrichment analysis performed using DESeq2 and Ingenuity Pathway Analysis (IPA, Qiagen). (a) Volcano plot of differential gene expression between keloid and non-keloid samples. (b) Bubble-volcano representation of selected pathways, where bubble size denotes the number of overlapping molecules. (c) Top canonical pathways ranked according to IPA activation *z*-score, and (d) Top canonical pathways ranked according to enrichment significance; orange bars indicate predicted activation and blue bars indicate predicted inhibition.

**Table 2:** Differential expression analysis results using DESeq2 for the consensus biomarkers derived using the multi-stage ML framework.

| Gene | Direction (ML) | $\log_2FC$ | Adjusted $p$-value |
| --- | --- | --- | --- |
| FASN | Up | -0.25 | 0.190 |
| CHN1 | Up | 1.52 | $4.4 \times 10^{-13}$ |
| GATA2 | Up | 1.31 | $2.3 \times 10^{-9}$ |
| KCNE4 | Up | 1.29 | $2.0 \times 10^{-8}$ |
| BGN | Up | 1.38 | $7.8 \times 10^{-8}$ |
| SDC4 | Down | -1.17 | $1.6 \times 10^{-9}$ |
| CLEC3B | Down | -1.86 | $3.9 \times 10^{-12}$ |
| TNXB | Down | -1.46 | $8.4 \times 10^{-9}$ |

### 3.7 Cell-Type-Specific Expression of Candidate Biomarkers

The harmonised single-cell atlas is shown in Figure 10. A comparison of the expression patterns of the eight consensus biomarkers across the annotated cell populations between cells originating from keloid and non-keloid samples is shown in Figure 11. The *log*_2_-fold change and multiple-testing-adjusted (p) values for the cell-type specific DGE analysis for the eight consensus biomarkers identified through the multi-stage ML framework are summarised in Figure 12.

**Figure 10:**
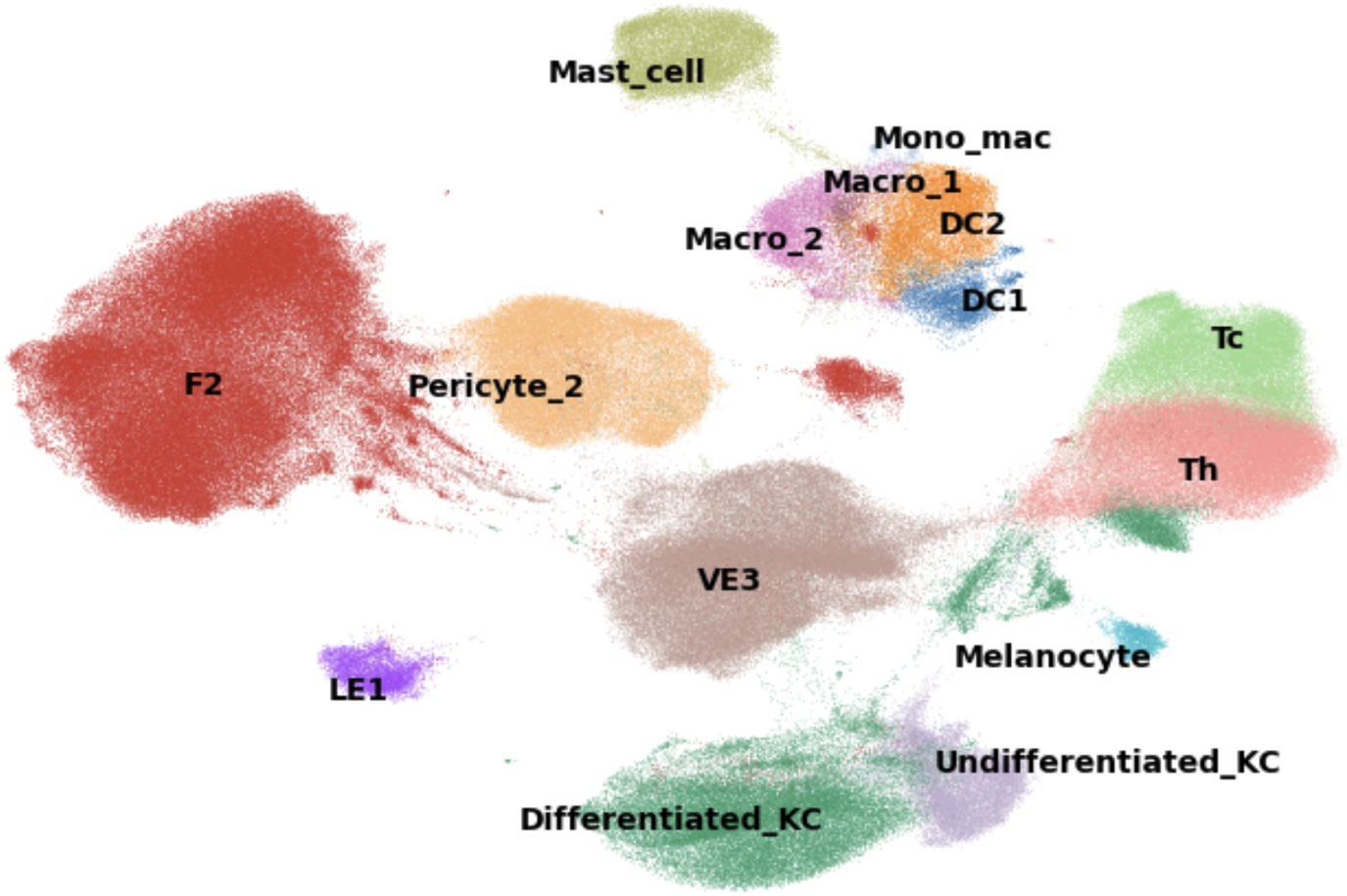
UMAP of the harmonised single-cell dataset. The identified clusters are colour-coded according to the cell-type annotations identified from CellTypist.

**Figure 11:**
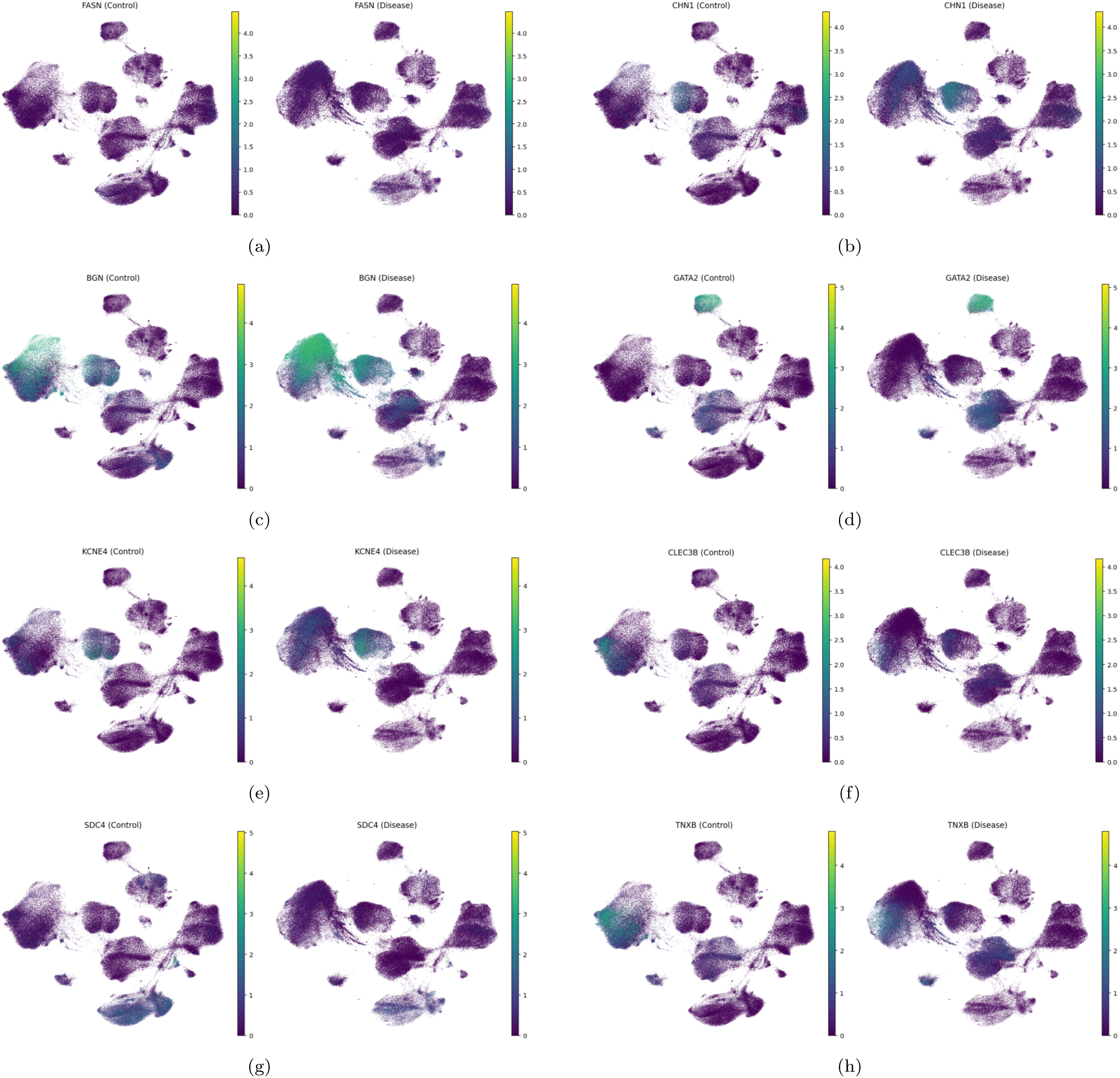
Single-cell expression patterns of the eight consensus biomarkers in non-keloid (control) and keloid (disease) samples. (a–e) Upregulated biomarkers; (f–h) downregulated biomarkers.

**Figure 12:**
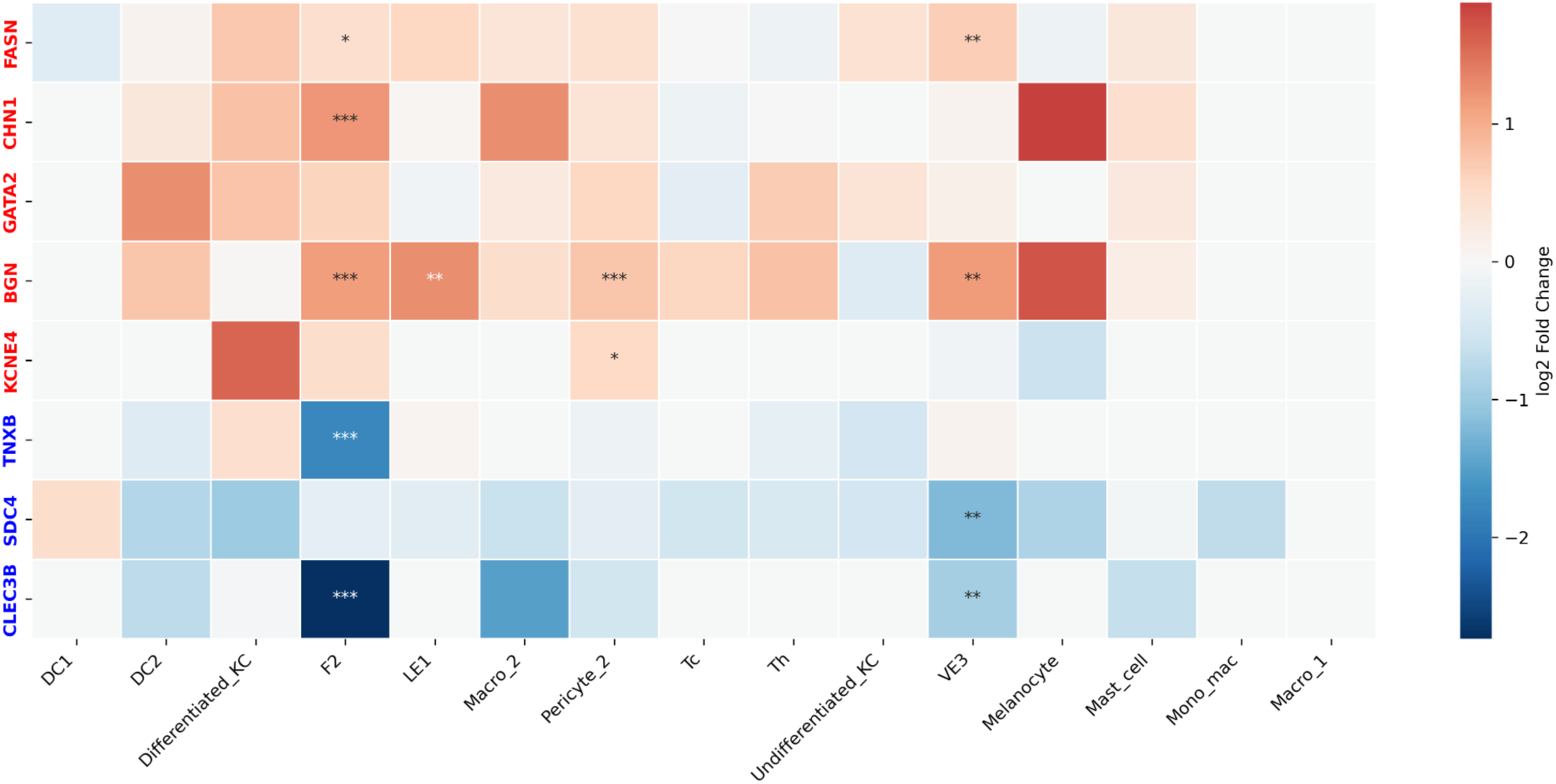
Cell-type-specific DGE of the eight consensus biomarkers identified through the multi-stage ML framework. Genes labelled in red and blue correspond to genes identified as upregulated and downregulated, respectively. Heatmap values correspond to DESeq2-derived log_2_ fold changes for keloid versus non-keloid samples within each annotated cell population. Asterisks denote adjusted *p*-value thresholds: * *p*_adj_ *<* 0.05, ** *p*_adj_ *<* 0.01, and *** *p*_adj_ *<* 0.001.

## 4 Discussion

In this study, we assembled, to the best of our knowledge, the largest curated cross-study transcriptomic cohort of keloid pathology currently available, integrating 81 samples from 13 independent studies across both bulk RNA sequencing and scRNA sequencing platforms and spanning four clinically relevant tissue classes: normal skin, normotrophic scar, hypertrophic scar, and keloid scar. By leveraging five complementary ML frameworks, we identified a set of robust and highly consistent biomarkers, summarised in Table 2, capable of distinguishing keloid tissue from other skin and scar phenotypes. Importantly, these biomarkers were selected through a multi-stage stability framework requiring consistency across multiple study-aware train-test partitionings, multiple ML classifiers, and bootstrap validation.

The employed classifiers rely on fundamentally different mathematical principles, and the feature-selection mechanisms is strongly related to the functional form of the ML algorithm making the selection. For the linear SVM, gene importance was inferred from the coefficients defining the maximum-margin separating hyperplanes. Similarly, in logistic regression, feature importance was determined from the regression coefficients governing the predicted class probabilities obtained from the softmax function. In contrast, random forests capture potentially complex non-linear interactions through ensembles of decision trees that iteratively split to maximize certain statistical measurements on the resulting partitions. Since tree models lack a closed form mathematical expression for the predicted label, feature importance and regulatory directionality were quantified using SHAP (SHapley Additive exPlanations) values, which estimate the contribution of each gene toward shifting model predictions toward or away from the keloid class. The convergence of these complementary ML frameworks toward a common set of biomarkers provides evidence that the identified genes capture robust transcriptomic signals that are not dependent on a particular modelling assumption, feature-selection strategy, or classifier functional form. Consequently, the selected biomarkers are more likely to represent biologically meaningful features of keloid pathology rather than artefacts of a specific framework.

Overall, all ML algorithms achieved reasonably strong classification performance, with mean multiclass accuracies and *F*_1_-scores exceeding 80%, while the binary classifiers, as expected, generally outperforming their multiclass counterparts due to the reduced complexity of the two-class discrimination problem. Interestingly, the multiclass random forest classifier did not substantially outperform the linear logistic regression and linear SVM models. This observation suggests that the transcriptomic separation between the skin and scar conditions can be captured effectively using approximately linear decision boundaries within the selected feature space, with limited additional predictive benefit obtained from modelling higher-order nonlinear interactions. The evaluation of alternative nonlinear ML frameworks on larger transcriptomic cohorts remains an avenue for future investigation.

The evaluation of the classifiers across multiple folds provided an empirical assessment of model robustness with respect to train-test partitioning. Despite the relatively limited cohort size (n=81), the distributions of the accuracy and *F*_1_-scores across the eligible folds (Figures 5 and 6) remained within a reasonable range for all classifiers, indicating relatively consistent predictive performance across independent allocations of studies to the training and testing sets. In general, the multiclass logistic regression and SVM classifiers demonstrated greater stability in the identification of important biomarkers than their binary counterparts, with the SVM yielding the largest number of consistently selected genes across folds. In contrast, the random forest classifier generalized poorly across the folds, as indicated by the poor convergence on the key biomarkers across the folds in Figure 7. Such a behaviour is not entirely surprising given the larger hyperparameter space, stochastic tree construction, and greater sensitivity to perturbations in the training data associated with random forest models, particularly in settings of high dimensional feature space, and further supports the notion that relatively simple and interpretable linear models may be sufficient to capture the dominant distinguishing transcriptomic signatures.

Inspection of the multiclass confusion matrices in Figure 3 reveals that the majority of misclassifications occurred between the scar phenotypes themselves, while normal skin samples were classified correctly in nearly all cases across the different ML frameworks. This finding is consistent with the exploratory PCA analysis (Figure 2(d)), in which normal skin samples exhibited clear separation from the scar classes. Together, these observations suggest that, in addition to achieving the classification objective of distinguishing the four tissue classes, the multiclass classifiers implicitly captured transcriptomic signatures associated with the presence of scarring. Furthermore, the few samples incorrectly classified as normal skin predominantly originated from the normotrophic scar class, which would be expected given that normal scars likely occupy an intermediate biological state between normal skin and pathological scarring, and therefore exhibit greater transcriptomic and molecular similarity to normal skin than the pathological scar phenotypes. Although the limited number of hypertrophic scar samples may have influenced some of the observed classification patterns, the overall structure of the misclassifications appears biologically plausible and suggests that the classifiers captured meaningful transcriptomic relationships among the different scar classes.

Finally, for interpretability purposes, the multiclass predictions were additionally collapsed into keloid versus non-keloid outcomes and compared against the corresponding binary classifiers, with the resulting sensitivity and specificity values summarised in Table S2. When comparing each binary classification framework (logistic regression or SVM) to its multiclass counterpart, the multiclass classifiers consistently exhibited higher specificity at the expense of reduced sensitivity. This finding supports the hypothesis that multiclass frameworks impose stricter transcriptomic discrimination between keloids and other scar phenotypes, thereby reducing false-positive keloid predictions arising from transcriptomic signatures associated with scarring more broadly. Conversely, in the binary setting, samples exhibiting transcriptomic features characteristic of pathological or mature scar tissue may be more readily assigned to the keloid class because the classifier is not explicitly required to distinguish between keloids, hypertrophic scars, and normotrophic scars, resulting in increased sensitivity but reduced specificity relative to their multiclass counterparts. Hence, binary and multiclass formulations capture complementary aspects of transcriptomics-driven keloid characterisation and further justify the multi-framework strategy adopted in this study for the extraction of robust biomarkers.

Having established the robustness of the biomarker selection framework, we next examine the biological significance of the identified genes. These biomarkers span pathways involved in ECM organisation, fibrosis, cellular adhesion, vascular biology, inflammation, and cellular metabolism, each of which has been implicated in wound healing and keloid pathogenesis. We begin with the downregulated biomarkers. *SDC4* encodes syndecan-4, a transmembrane heparan sulfate proteoglycan that couples the ECM to the actin cytoskeleton, thereby playing a central role in mechan-otransduction, focal adhesion formation and fibroblast migration. [22] *CLEC3B* is a plasminogen-binding protein that facilitates the activation of plasminogen into plasmin, a key mediator of fibrinolysis (the removal of blood clots) and ECM turnover. Through the activation of plasminogen, *CLEC3B* contributes to degradation of fibrin and non-collagenous ECM components, including laminin and fibronectin, thereby promoting tissue remodelling and wound resolution. [23] Furthermore, *CLEC3B* has been associated with increased activity of matrix metalloproteinases, including MMP-2 and MMP-9, which contribute to the degradation of excess type I and type III collagen. [24] A recent work has further demonstrated a direct antifibrotic role for *CLEC3B* ; overexpression of *CLEC3B* attenuated skin fibrosis in bleomycin-induced fibrosis models and reduced fibrosis severity in keloid xenografts, while knockdown activated the Wnt/(β)-catenin pathway and promoted profibrotic signalling. [25] Finally, *TNXB* encodes the ECM glycoprotein tenascin-X, a key organiser of collagen fibril architecture into ordered and functional ECM networks. The anti-fibrotic properties of tenascin-X have been attributed to its anti-adhesive properties, [26] which affect the ECM mediated fibroblast migration, as well as due to its role in regulating TGFβ, a key signalling factor in fibrosis and wound healing. [27]

Keloids are conventionally interpreted through the lens of excessive collagen deposition, fibroblast activation, and inflammatory signalling; these mechanisms are indeed correlated with the transcriptomic biomarkers that are pronounced in the present dataset (Figure 7, particularly genes identified from multiclass SVM, and Table S1), along with the inferred upregulated signalling pathways related to collagen deposition and ECM components from the IPA analysis in Figure 9(c). Interestingly, however, the downregulation of these genes collectively suggests that an additional contributor to keloid pathogenesis and fibrotic persistence in the lesion microenvironment may be the loss of physiological mechanisms responsible for ECM homeostasis and wound resolution. Specifically, the coordinated suppression of *SDC4*, *CLEC3B*, and *TNXB* points toward impaired matrix turnover, altered cell-matrix mechanotransduction, and diminished antifibrotic regulation. The notion of impaired homeostatic regulation is further supported by the IPA analysis (Figure 9(d)), which identified nonsense-mediated decay and EIF2 signalling, the quality-control pathways for mRNAs and proteins, respectively, as among the most significantly disrupted (downregulated) pathways.

We next consider the upregulated biomarkers. *BGN* encodes biglycan, a small leucine-rich ECM proteoglycan that plays multiple roles during wound healing and tissue remodelling. During the inflammatory and proliferative phases of healing, biglycan participates in cellular signalling between the ECM and the skin cells. Furthermore, biglycan promotes angiogenesis during tissue repair, [28] binds collagen fibrils and regulates their assembly while increasing fibril diameter, [29] and has been implicated in he differentiation of fibroblasts toward a proto-myofibroblast phenotype, [30] as well as in epithelial-mesenchymal transition (EMT), [31] an important mechanism that is recgonised to give rise to the invasive and fibroproliferative characteristics of keloid pathology. [32] *KCNE4*, which encodes a potassium-channel regulatory subunit, has been implicated in the regulation of arterial tone and vascular reactivity. Similarly, *GATA2* is a transcription factor with well-established roles in endothelial-cell development, vascular maintenance, and angiogenic responses. For *CHN1*, altough its is classically associated with neuronal development and axonal guidance, recent studies have demonstrated a direct role for *CHN1* in promoting EMT in several cancer types, [33] which, as previously mentioned has a recognised role in keloid pathogenesis. [32] Interestingly, the apparent vascular signature suggested by *KCNE4*, *BGN*, and *GATA2* is further supported by the cell-type-specific differential expression analysis in Section 3.7: while significant differential expression of several biomarkers was observed in fibroblast populations, as expected, robust and statistically significant expression differences were also detected in vascular endothelial cells (vE3) which line the vessel walls and regulate vasofilation via shear-stress mediated signalling, with additional contributions from pericytes, which regulate capillary diameter and microvascular function, as well as lymphatic endothelial cells, thereby implicating vascular remodelling and functionality in the development and persistence of keloid lesions.

All the genes discussed above exhibited strong statistical evidence of differential expression (*p <* 0.001) between the keloid and non-keloid samples in the DGE analysis, as shown in Table 2, with directionality consistent with that identified by the ML framework. One notable exception was *FASN*, which emerged as a highly consistent biomarker across the ML analyses despite demonstrating weak and non-significant downregulation in the DGE analysis. Specifically, *FASN* exhibited consistently positive feature importance across the ML classifiers and permissible fold partitions, with the corresponding bootstrap binary logistic regression 95% confidence interval remaining entirely above zero. In contrast, the DGE analysis suggested a small, non-significant negative (log_2_) fold change between the keloid and non-keloid cohorts (Table 2). Examination of the cell-type-specific differential expression patterns in Figure 12 revealed a general trend toward elevated *FASN* expression in keloid-derived cells, with statistically significant upregulation observed in fibroblast (F2) and vascular endothelial (vE3) populations. These scRNA cell type-specific findings are therefore more consistent with the directionality inferred from the ML analysis than from the bulk DGE comparison. One possible explanation for this discrepancy is that DGE and ML frameworks quantify fundamentally different properties of the data: DGE analysis evaluates differences in average expression between groups, whereas ML-based feature selection identifies genes that contribute most strongly to classification performance. The genes that provide strong discriminatory information may not necessarily exhibit the largest average fold changes. This is particularly true in heterogeneous datasets composed of multiple cell populations, studies, and sequencing platforms, where biologically informative signals may be diluted when averaged across samples yet remain sufficiently informative for classification algorithms to repeatedly select them as predictive features.

Accordingly, *FASN* may represent a biologically relevant component of keloid pathology that would have been overlooked through reliance on DGE analysis alone. *FASN* encodes fatty acid synthase, a multifunctional enzyme responsible for de novo lipogenesis, through which long-chain saturated fatty acids, primarily palmitate, are synthesized from acetyl-CoA and malonyl-CoA using NADPH. Interestingly, *FASN* is among the most consistently overexpressed metabolic enzymes across multiple solid tumours, including breast, prostate, pancreatic, and ovarian cancers, where increased fatty-acid synthesis supports membrane biogenesis, energy storage, and the anabolic demands of rapidly proliferating cells. [34] In cancer biology, elevated *FASN* expression is frequently discussed within the broader context of Warburg effect, [35] in which cancer cells consume glucose and convert it into lactate via anaerobic respiration for the rapid glycolytic carbon flux of acetyl-coA required for lipogenesis, along with the rapid, albeit inefficient, generation usable energy (ATP). Furthermore, elevanted FASN levels in tumours have been shown to confer apoptotic and chemotherapy resistance in some tumours. [36] The identification of *FASN* is particularly intriguing given the tumour-like features of keloids, including persistent proliferation, apoptotic resistance and invasion of surrounding healthy skin. This notion of altered metabolic programming in keloids is in line with recent studies that highlight metabolic pathways and amino acid biosynthesis as key affected pathways identified by functional enrichment analysis, [37] and are further supported by the results of the IPA in Section 3.6. For instance, the IPA analysis in Figure 9(d) identifies the ‘response of *GCN2* to amino acid deficiency’ as a significantly downregulated pathway, which suggests that cells fail to properly suppress protein translation during starvation, and continue to consume amino acid pools instead of engaging the normal metabolic stress response.

The particular role of altered lipid metabolism in keloid formation and progression has received increasing attention in recent years. [38–40] However, the lipid landscape in keloids appears complex, with different lipid species exhibiting distinct patterns of expression. For example, arachidonic acid and several polyunsaturated fatty acids have been reported to be elevated in keloid tissue, whereas short-chain fatty acids such as butyrate and valerate, as well as certain essential fatty acids including α-linolenic acid, have been found to be reduced. [39, 40] These findings suggest that keloid pathogenesis may involve broader metabolic reprogramming rather than a simple increase or decrease in lipid synthesis. Of particular interest is the work of Yan et al., [41] who reported lower overall FASN mRNA and protein expression in keloid tissue compared with normal scar tissue, yet significantly higher FASN mRNA expression in keloid-derived fibroblasts when fibroblasts were analysed in isolation, while fibroblast FASN protein levels did not differ significantly between the two groups. In the present study, FASN was significantly upregulated in fibroblasts (F2) and vascular endothelial cells (vE3) (Figure 12), while exhibiting non-significant trends toward downregulation in several other cell populations. These findings may help further reconcile the apparent discrepancy between the ML and DGE analyses. While overall FASN expression across whole tissue samples may be modestly reduced, its expression appears concentrated within specific cell populations central to keloid pathology, particularly fibroblasts and endothelial cells, which may be key drivers of the metabolic reprogramming observed in keloids. Together, these observations highlight the importance of considering cell-type-specific expression patterns when interpreting bulk tissue analyses and suggest that further functional studies are required to clarify the mechanistic contribution of FASN to keloid progression.

Finally, analysis of the GSEA results in Figure S1 reveals enhanced communication between cells and their extracellular microenvironment in keloid lesions, as evidenced by the enrichment of glycan and glycosaminoglycan biosynthesis pathways. N-glycans are important post-translational modifications of cell-surface receptors, including integrins and growth factor receptors, and increased N-glycan biosynthesis can increase cell-ECM signalling and cellular responsiveness to extracellular cues. [42] Similarly, glycosaminoglycans are major ECM components that regulate cellular communication by binding growth factors, cytokines, and other signalling molecules. [43]

In a high-dimensional transcriptomic dataset, not all features are informative, and the inclusion of irrelevant or highly correlated genes may even impair the learning process by introducing noise and increasing estimator variance. Accordingly, our framework restricted the analysis to a subset of features selected independently within each outer fold, as described in Section 5.4, by incorporating the feature-selection hyperparameters *K* and *F_thresh_* into the cross-validation grid search. Indeed, the hyperparameter optimisation results consistently indicated that all ML algorithms generally benefited from reduced feature space (lower values of *K*), suggesting that increasing the number of included genes does not necessarily improve predictive accuracy and may instead introduce additional noise into the feature space. Future work could explore more sophisticated feature-selection approaches, such as recursive feature elimination, in which less informative genes are iteratively removed based on their contribution to model performance.

Following evaluation of the ML algorithms across the three outer folds, the feature-selection hyperparameters for each classifier were fixed to the combination yielding the highest mean performance across the outer folds and subsequently used throughout the partition-stability analysis. While fixing these parameters may underestimate the full variability associated with repeated feature-selection optimisation across all eligible folds, performing a complete grid search over the feature-selection hyperparameter space for each of the 56 permissible partition configurations would be computationally prohibitive. Furthermore, excessive hyperparameter optimisation may itself introduce an additional source of variability and overfitting to particular cross-validation configurations. [44] Importantly, despite fixing the feature-selection hyperparameters, the classifiers maintained strong and relatively stable predictive performance across the permissible folds (Figures 5 and 6).

As described in Section 5.2, the harmonization procedure used to mitigate study, chemistry, and platform-specific effects was performed prior to train-test partitioning and therefore introduces a limited degree of information leakage, since the corrected representation of each sample is influenced by the global structure of the dataset. Nevertheless, comparison of Figures 1 and 2 demonstrates that such harmonization was essential for reducing dominant technical sources of variation while preserving biologically meaningful separation between the skin and scar phenotypes. Without this correction step, the ML algorithms would be at substantial risk of learning study, platform, or chemistry-specific artefacts rather than transcriptomic signatures associated with scar pathology. Importantly, the primary objective of this study was not the development of a universally deployable classifier capable of annotating future external samples, but rather the use of ML as a tool for biomarker discovery across the largest curated collection of transcriptomic datasets currently available. Within this context, the benefits of reducing technical confounding are likely to outweigh the limited information sharing introduced by the harmonization procedure. Furthermore, this harmonization step represents the only stage of the workflow involving dataset-wide information; all subsequent stages of the analysis, including feature selection, z-score standardization, model fitting, biomarker extraction, and hyperparameter optimization, were performed under strict train-test separation, ensuring that no information from the held-out testing partitions was used during model training.

Several other limitations of the present study should be acknowledged. First, the analysis necessarily relied on the clinical annotations and sample classifications reported by the original studies, and the distinction between scar phenotypes may not have been entirely consistent across studies. Second, the cell-type-specific analyses relied on CellTypist annotations [45] derived from reference atlases of healthy human skin. [46] While this approach provides a consistent annotation framework across studies, diseased tissues such as keloids may contain altered cellular states, recognized transitional populations in fibroblast, macrophage and endothelial cell populations, and pathological differentiation programs that are not fully represented within healthy reference datasets. Finally, the multiclass classification framework treated the four tissue classes as independent categorical labels. However, the classification results and confusion matrices suggest that biologically meaningful relationships exist between the classes, as expected with normal skin, normotrophic scar, hypertrophic scar, and keloid scar potentially representing progressively distinct stages along a spectrum of wound-healing and fibrotic remodelling. Future studies could therefore investigate explicitly encoding relative similarities between tissue classes, for instance, by using numbered categories.

In conclusion, the robust biomarkers identified in this study provide insight into the molecular mechanisms underlying keloid initiation and progression, which may ultimately facilitate the development of more targeted therapeutic interventions. The translation of these findings into specific treatment strategies is beyond the scope of the present study; nevertheless, we provide two simple examples illustrating how the identified biomarkers may potentially inform future therapeutic development. First, *CLEC3B* was consistently identified as a downregulated biomarker and has recently been shown to attenuate fibrosis and reduce keloid severity when administered as recombinant protein in experimental fibrosis and keloid xenograft models. [25] Given its role in plasmin-mediated ECM turnover and wound resolution, restoration of *CLEC3B* signalling through recombinant protein delivery, or alternative approaches aimed at enhancing the downstream plasminogen activation such as the recombinant plasminogen activators used in thrombolytic therapy, may represent a potential avenue for future antifibrotic local intervention. Furthermore, *FASN* is a well-established metabolic enzyme and druggable target that is overexpressed in numerous human malignancies, for which several pharmacological inhibitors, including TVB-2640 (denifanstat), TVB-3664, and experimental compounds such as C75 and cerulenin, have been developed and evaluated in preclinical and clinical settings. [47, 48]

Since *FASN* has shown consistent upregulation in our study, the local administration of FASN inhibitors could in principle be another treatment strategy for keloid lesions. Of course, such applications remain to be investigated experimentally.

## 5 Materials and Methods

### 5.1 Dataset Assembly and Curation

Transcriptomic datasets were identified through a systematic search of the Gene Expression Omnibus (GEO) repository [49] and related public databases for both bulk RNA-seq and scRNA-seq studies of human skin samples, corresponding to one of four conditions of interest: keloid scar, hypertrophic scar, normotrophic scar, or normal skin. With the exception of one bulk RNA dataset (OEP00002674), [16] which was obtained from the National Omics Data Encyclopedia, [50] the remaining samples can be retreived from the GEO database. Several categories of datasets were excluded. These included microarray studies (for instance, using Affymetrix technology), datasets containing only spatial transcriptomic measurements despite being incorrectly annotated as scRNA-seq datasets in public repositories, [9], datasets containing transcriptomic measurements from isolated cells rather than intact tissue biopsies (see Section 2), studies in which samples were subjected to experimental treatments (*e.g.*, therapeutic injections), and samples associated with comorbid conditions such as systemic sclerosis. [51] In addition, datasets containing biopsies maintained in culture for multiple days were excluded because prolonged exposure to culture conditions may substantially alter gene expression profiles and cause genetic drifts. [52, 53] Datasets lacking raw count information and containing only transformed or normalized expression values were also excluded.

Several datasets required additional curation. For instance, for the GSE220300 dataset, [54] active and inactive keloids were merged into a single keloid category, while normal scars and mature scars were grouped into a single normal scar category. Furthermore, central and peripheral biopsies originating from the same keloid lesion were aggregated to generate a single patient-level sample. For the GSE181316 dataset, [55] bilateral keloid biopsies obtained from the same patient were retained as independent samples. Similarly, biopsies corresponding to both hypertrophic scar and keloid tissue from the same patient in the GSE243716 dataset [7] were treated as separate samples. While samples originating from the same patient may share patient-specific biological characteristics that might bias the ML framework or introduce leakage upon partitioning, the downstream partitioning and cross-validation procedures were performed at the study level (Section 5.3), rather than at the sample level, thereby preventing information leakage from samples of the same origin. Following quality control and curation, the final harmonized dataset inlcuded 81 samples collected across 13 independent studies, comprising 59 pseudo-bulked scRNA-seq samples and 22 bulk RNA-seq samples. The resulting study composition and sample metadata are summarized in Table 3.

**Table 3:** Summary of transcriptomic datasets included in the final harmonized cohort. The *Chemistry* column indicates the 10x Genomics scRNA sequencing chemistry used for the scRNA datasets.

| Study | Platform | Chemistry | Keloid | Hypertrophic | Normal Scar | Normal Skin | Total |
| --- | --- | --- | --- | --- | --- | --- | --- |
| GSE156326 [19] | scRNA-seq | 10x v2/v3 | 0 | 3 | 0 | 3 | 6 |
| GSE163973 [14] | scRNA-seq | 10x v2 | 3 | 0 | 3 | 0 | 6 |
| GSE181297 [13] | scRNA-seq | 10x v2 | 2 | 0 | 1 | 0 | 3 |
| GSE243716 [7] | scRNA-seq | 10x v3 | 1 | 1 | 0 | 0 | 2 |
| GSE266334 [11] | scRNA-seq | 10x v3 | 3 | 0 | 0 | 1 | 4 |
| GSE270438 [15] | scRNA-seq | 10x v3 | 3 | 0 | 3 | 0 | 6 |
| GSE303592 [10] | scRNA-seq | 10x v3 | 1 | 0 | 0 | 1 | 2 |
| GSE188952 [12] | bulk RNA-seq | N/A | 4 | 5 | 3 | 0 | 12 |
| GSE181316 [55] | scRNA-seq | 10x v3 | 4 | 0 | 3 | 1 | 8 |
| OEP0002674 [16] | bulk RNA-seq | N/A | 0 | 5 | 0 | 5 | 10 |
| GSE130973 [56] | scRNA-seq | 10x v2 | 0 | 0 | 0 | 5 | 5 |
| GSE138669 [51] | scRNA-seq | 10x v1/v2 | 0 | 0 | 0 | 10 | 10 |
| GSE220300 [54] | scRNA-seq | 10x v2 | 4 | 0 | 3 | 0 | 7 |
| **Total** |  |  | **25** | **14** | **16** | **26** | **81** |

### 5.2 Data Preprocessing and Harmonization

The datasets analysed in this study were generated using two distinct high-throughput transcriptomic sequencing technologies that differ in resolution, sequencing depth, and output structure. Specifically, scRNA-seq quantifies gene expression levels for each individual cell *c*, producing an expression matrix for each sample *s* of the form **M***^s^* ∈ *R^C,G^* where *G* denotes the number of genes. In contrast, bulk RNA sequencing measures average gene expression across a whole tissue sample, producing a single expression vector *M^s^* ∈ *R^G^*. To enable direct comparison between transcriptomic profiles generated from these two platforms, it was necessary to standardise their expression formats. Accordingly, the following preprocessing steps were applied to aggregate the scRNA-seq data into a pseudo-bulk representation comparable to bulk RNA-seq measurements:

1. Cells were filtered according to standard quality-control criteria based on mitochondrial gene expression (*g_m_*), total transcript count, and the number of detected genes per cell:

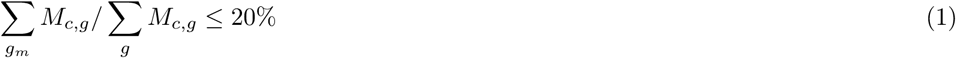

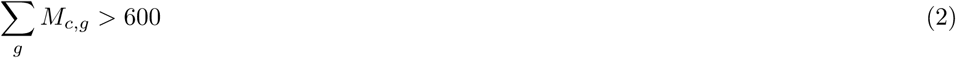

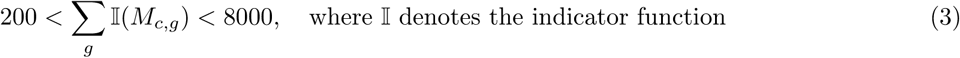

2. Genes detected in fewer than five cells after cell-level filtering were removed 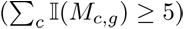.

3. For each sample, raw gene-expression counts were summed across all remaining cells to generate a pseudo-bulk expression vector M^(*s*)^ ∈ ℝ*^G^*.

To assemble the gene expression vectors (M^(*s*)^) across all samples *s* presented in Table 3, gene nomenclature was standardised by converting bulk RNA-seq Ensembl identifiers to gene symbols with approximately 85% gene retention. Subsequently, only genes shared across all datasets were retained for downstream analysis. To account for differences in sequencing depth and library size across samples, each sample was independently normalised using log 2-counts-per-million (log2-CPM) normalisation; importantly, the normalisation was performed independently for each sample to minimise information leakage during downstream ML analyses.

We first performed an exploratory analysis to assess the extent to which variation in gene expression across samples could be attributed to biological versus technical sources. Principal component analysis (PCA) was applied to the combined log-normalized expression matrix to visualize global sample organization, identify potential batch effects, and evaluate condition-driven separation. The samples were projected onto the first three principal components, corresponding to the largest orthogonal axes of variation. Given the absence of discernible separation between v3.0 and v3.1 samples in the exploratory PCA, the two scRNA-seq chemistries were merged into a single category in order to reduce the number of technical covariates included in the subsequent batch-correction and harmonization procedures.

The dominant sources of variation in gene expression across the samples can be attributed to technical factors, which are: sequencing platform (2 categories: scRNA-seq versus bulk RNA-seq), experimental source (13 studies, or categories), and scRNA-seq library preparation chemistry/protocol (3 categories). Applying any ML models to the uncorrected expression matrix would result in the model learning platform and batch-specific technical artefacts rather than biologically meaningful and distinguishing transcriptomic features. To correct for such technical sources of variation, we employed ComBat, [57] a statistical harmonisation algorithm in R based on an empirical Bayes framework, which estimates and removes unwanted technical variation while attempting to preserve the biologically relevant signal. In essence, ComBat assumes a linear regression model in which the observed expression of gene *g* in sample *i* is decomposed into four components, as shown in Equation 4: (1) the baseline expression of the gene α*_g_*, (2) the biological contribution of the covariates of interest (e.g., scar type) *X_i_*β*_g_*, (3) an additive batch-specific effect γ*_g,b_*_(*i*)_, and (4) a multiplicative batch-specific scaling factor δ*_g,b_*_(*i*)_:

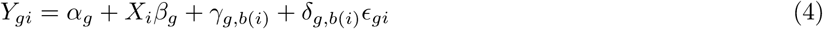

where *b*(*i*) denotes the batch assignment of sample *i*, specified through a user-defined batch annotation.

Given the relatively small cohort size (n=81), modelling these technical factors independently would substantially increase the size of the covariance correction matrix and risk overcorrection or unstable parameter estimation. However, these technical covariates exhibited a high degree of overlap and dependency. Specifically, platform variation was inherently nested within study variation, since samples originating from the same study were generated using a either one of the two sequencing platform. Furthermore, with the exception of two scRNA-seq studies, samples within the same scRNA-seq study were processed using the same library preparation chemistry.

Consequently, rather than modelling platform, study, and chemistry as independent batch variables, we constructed a composite technical covariate combining study identity and, where applicable, chemistry information for studies containing multiple chemistries. This resulted in 15 unique batch combinations capturing the cumulative additive and multiplicative technical effects of platform, study, and scRNA chemistry, as captured in γ*_g,b_*_(*i*)_ and δ*_g,b_*_(*i*)_ in Equation 4, while maintaining a parsimonious batch structure appropriate for the available sample size. We note that, as indicated by Equation 4, the transformation applied to the log-normalized expression value of a given gene in a particular sample depends collectively on the log-normalized expression distribution of that gene across all samples. As such, the use of ComBat for dataset harmonisation introduces some limited degree of information leakage into the downstream ML analyses, since following test-train sample partitioning, the transformed feature representation of samples in the training set is influenced, to some extent, by information originating from the testing set. The implications of such data leakage on the ML classification and the corresponding derived biomarkers are discussed in Section 4.

### 5.3 Sample Stratification

We implemented a study-aware stratified (k)-fold cross-validation strategy in which all samples originating from the same experiment were kept together within a single partition to ensure that the models are evaluated on completely unseen experimental batches and eliminate any study-specific information leakage between training and testing partitions. In addition, the partitioning was constrained to ensure similar distributions of skin categories in the training and testing sets. Specifically, the test partition was required to contain between 21 and 27 samples (out of 81) and include at least three samples from each class. One notable problem is that partitioning an already limited cohort (n=81) into fixed training and testing sets may reduce the amount of information available for model training and keep useful examples from training, even if the testing and training sets share similar categorical distributions. This issue is particularly exacerbated in high-dimensional transcriptomic datasets where the number of features greatly exceeds the number of samples, potentially increasing estimator variance and reducing the stability of the identified biomarkers. Hence, the train-test partitioning was repeated across multiple outer folds with mutually exclusive testing sets to maximize the use of the available samples despite the limited cohort size (n=81), allowing all samples to contribute to model training and hyperparameter tuning across the collection of folds and reducing the effect of the partition choice.

One particular limitation was the low availability of hypertrophic scar samples, which were present in only four experiments, one of which contained a single sample. Consequently, the three experiments containing more than one hypertrophic scar sample were designated as anchor studies, with one anchor assigned to the testing partition of each outer fold. As a result, the maximum feasible number of folds was restricted to three. The remaining experimental batches were then automatically distributed across the three outer folds under the above constraints to achieve balanced class representation while preserving complete study segregation between training and testing partitions. Based on these criteria, we were able to construct 56 unique folds that could be assembled into 52 valid train-test partition combinations, each comprised of three outer folds. The first partition configuration was used for tuning the model hyperparameters, while the remaining fold combinations were utilised for partition stability analysis (see Section 5.5). For each of the three outer cross-validation folds, model hyperparameters were tuned exclusively within the corresponding training partition using study-aware inner cross-validation.

### 5.4 Transcriptomic Feature Selection

With approximately 8000 genes remaining in the harmonised dataset after filtering, the dimensionality of the feature space is large relative to the available sample size (n=81), increasing the risk of noise-driven classification and overfitting. To address this, feature screening and reduction were performed independently within each training outer fold in order to identify a preliminary subset of biologically distinguishing genes while preventing information leakage from the testing partitions. Specifically, the feature selection procedure was governed by two parameters, *K* and *F_thresh_*. For each skin/scar condition and for each gene, the *log*_2_ fold change between the condition of interest and the remaining samples (“condition versus rest”) was estimated from the difference in the average corrected log2-CPM values. Statistical significance was then assessed using a two-sided t-test, and the Benjamini Hochberg procedure was applied with a false discovery rate threshold of *F_thresh_* to account for multiple hypothesis testing. From the subset of genes for which the null hypothesis was rejected, the top *K* genes with the highest absolute *log*_2_ fold changes for the corresponding condition were selected. The final feature set for a given fold was then obtained by combining the selected genes across all conditions. To ensure that feature optimisation was not influenced by the held-out testing partitions (*i*.*e*., no data snooping was introduced), these two parameters were treated as hyperparameters of the feature selection pipeline and tuned exclusively within the training sets of the three folds, alongside the parameters specific to the ML algorithm (see Section 5.5), using study-aware cross-validation, as descibed in Section 5.3. A total of 390 pairwise combinations of the feature selection hyperparameters were investigated:

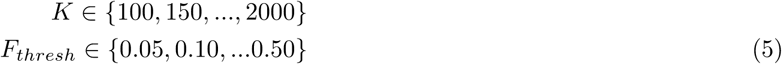

### 5.5 Machine Learning Algorithms

Following feature selection, the training data within each of the three outer folds were z-standardised to ensure comparable scaling across genes and prevent high-variance features from disproportionately influencing model training. The resulting transformation was then applied to the corresponding testing partition in each outer fold. Next, we tried multiple ML frameworks that utilize distinct classification algorithms: (1) multiclass logistic regression (one-vs-rest), (2) multiclass support vector machines (SVMs) with linear kernels, (3) binary logistic regression, (4) binary linear SVMs, and (5) multiclass random forest classifiers. Because different algorithms rely on distinct paradigms and functional forms to construct decision boundaries and determine feature importance, leveraging classifier algorithms reduces the risk of biasing biomarker discovery toward model-specific patterns, artefacts or methodological limitations.

In particular, the use of both binary and multiclass classification frameworks was intended to capture complementary aspects of transcriptomic separation. Binary models identify transcriptomic features broadly associated with keloid pathology, favouring higher sensitivity to general keloid-related molecular signals. In contrast, multiclass models can favour higher specificity by distinguishing keloids from individual skin and scar conditions that share overlapping characteristics of tissue remodelling, such as the generalised scarring and inflammatory responses observed in normal and hypertrophic scars, as well as increased collagen deposition in hypertrophic scars. Consensus biomarkers found at the intersection of these complementary classifying frameworks can represent highly refined, robust signature genes that are uniquely characteristic of keloid scarring pathology.

The hyperparameter search included both the feature-selection parameters *K* and *F_thresh_* described in Section 5.4, as well as the algorithm-specific hyperparameters associated with each ML classifier. For logistic regression, models were trained using the saga solver with elastic-net regularisation, introducing two algorithm-specific hyperparameters to tune: the regularisation strength *C* and the elastic-net mixing parameter *l*1*_ratio_*, controlling the balance between ridge (*L*_2_) and lasso (*L*_1_) penalties. For support vector machines (SVMs), a linear kernel was adopted, introducing a single regularisation parameter *C* controlling the tradeoff between classification performance and margin maximisation. For random forest classifiers, hyperparameter optimisation included the number of trees, maximum tree depth, minimum samples required for node splitting, minimum samples per leaf, and the number of features considered at each split.

All classifiers were optimised using the macro-averaged *F*_1_-score for multiclass classification and the standard binary *F*_1_-score for binary classification. These metrics were selected because they are less sensitive to class imbalance than overall accuracy, thereby preventing dominant classes from disproportionately influencing model optimisation and evaluation. The optimal hyperparameter combination identified during inner cross-validation was subsequently used to refit the model on the corresponding full outer training partition within each outer fold, after which predictive performance was evaluated on the corresponding held-out testing partition.

### 5.6 Partition Stability Analysis

A clear variability in classification performance has been observed across the outer folds in Figures 3 and 4, as also indicated by the standard deviations in Table 1, with outer fold 1 generally outperforming folds 2 and 3. This suggests that classification performance, and potentially the corresponding biomarkers identified by the ML frameworks, may exhibit sensitivity to the particular study-aware partitioning adopted, especially given the limited cohort size (*n* = 81). To this end, we subsequently employ a stability analysis to evaluate classification performance across all 56 outer-fold partition combinations satisfying the criteria outlined in Section 5.2, and to identify biomarkers demonstrating consistency across both folds and ML classifiers.

For the stability analysis, the feature-selection hyperparameters previously optimized for each ML algorithm using the initial partition configuration were fixed at the values yielding the highest mean outer-fold performance across the three outer folds of that configuration. This was done to avoid the substantial computational burden associated with repeatedly optimizing the high-dimensional feature-selection search space across all 56 eligible partition configurations. The selected genes were recomputed independently within each training partition using the fixed feature-selection hyperparameters, and the remaining classifier-specific hyperparameters were subsequently tuned independently within each outer fold using study-aware inner cross-validation, as described previously.

### 5.7 Biomarker Stability Analysis

To identify the stable transcriptomic biomarkers across the 56 eligible folds, We extracted and quantified feature importance and regulatory directionality (upregulation/downregulation) from the trained models for each ML algorithm. For each fold, up to 200 upregulated and 200 downregulated genes were selected. For logistic regression and linear SVM classifiers, gene ranking was based on the magnitude and sign of the learned coefficients arising from the linear decision functions, with the largest positive and negative coefficients corresponding to the most strongly upregulated and downregulated keloid-associated features, respectively. For random forest classifiers, since there is no explicit mathematical formulation for the output, feature importance and regulatory directionality were quantified using SHAP (SHapley Additive exPlanations) values, [58] which estimate the contribution of each gene toward shifting model predictions toward or away from the keloid class. Stable biomarkers for each ML algorithm were subsequently defined as genes exhibiting consistent upregulation or downregulation across more than 70% of the eligible outer-fold partitionings. Finally, to identify a parsimonious set of transcriptomic biomarkers with the highest discriminatory power, we restricted our analysis to stable genes identified by at least three of the five ML frameworks. As discussed in Section 2, this consensus requirement was introduced to prioritize biomarkers exhibiting robustness across distinct classification paradigms and feature-ranking methodologies, thereby reducing dependence on any single model architecture.

### 5.8 Bootstrap Validation of Candidate Biomarkers

These genes identified from Section 5.7 were then used as features in a final step using a binary logistic regression model fitted on the full harmonized dataset. While previous analyses focused on model evaluation in tandem with feature importance ranking, no train-test partitioning was performed at this final stage because the objective was the final characterisation of the most robust transcriptomic biomarkers using all available samples and the consensus feature set identified across the multiple ML frameworks. A study-aware group CV procedure was also used to tune the regularization parameter *C* (while an *L*_2_ penalty was adopted to avoid forcing coefficients of the reduced biomarker panel to exactly zero).

To quantify the uncertainty in the estimated biomarker effects, the final fitting step was repeated across 1000 bootstrap simulations, with each bootstrap dataset generated by independently sampling the original cohort of 81 samples with replacement to generate the regression coefficients and their corresponding confidence intervals. To determine whether the sign of each coefficient for the 14 genes remained significantly aligned with the direction of regulation inferred from the preceding stability analysis, one-sided empirical (p)-values were computed for the coefficients across the bootstrap simulations. The genes whose bootstrap-validation coefficients exhibited directionality consistent with that identified in the biomarker stability analysis (Section 5.7) demonstrated remarkable consistency across study-aware partitioning, repeated stability analyses, multiple ML classifiers, and bootstrap resampling. Collectively, these successive lines of evidence provide strong support for the robustness of these genes as discriminatory transcriptomic biomarkers of keloid pathology.

### 5.9 Differential Gene Expression Analysis

We compare the results obtained from the multi-stage ML framework with those from a traditional DGE analysis of the original bulk RNA-seq and pseudo-bulked scRNA-seq samples. Using DESeq2, [59] we conduct a binary keloid versus non-keloid comparison while accounting for the composite study/chemistry batch effect and preserving the biological condition of interest, and extract resulting *log*_2_ fold changes and multiple-testing-adjusted (p)-values for the eight consensus biomarkers. In addition, the DGE results for several canonical fibrosis- and keloid-associated genes frequently reported in the literature, including members of the transforming growth factor beta *TGF* β signalling pathway, collagen genes, ECM proteins, and wound-healing markers commonly implicated in keloid pathogenesis, are provided in Supplementary Table S1. The DESeq2-derived *log*_2_ fold changes and adjusted *p*-values were imported into Ingenuity Pathway Analysis (IPA, Qiagen) [60] to identify signalling, metabolic, and cellular pathways potentially activated or inhibited in keloid pathology.

### 5.10 Cell-Type-Specific Characterization of Candidate Biomarkers

Following the pathway-level analyses described above, we next investigated whether the eight consensus biomarkers identified through the ML stability analysis exhibited cell-type-specific expression patterns within the scRNA-seq datasets. To this end, all available scRNA-seq samples (21 keloid and 38 non-keloid samples) were reprocessed and integrated into a unified single-cell atlas. Standard cell-level quality-control filtering was first applied independently to each sample as described in Section 5.2. The filtered datasets were subsequently merged using an outer-join strategy, assigning zero counts to missing observations, yielding an integrated dataset comprising approximately 4 × 10^5^ cells. Genes detected in fewer than ten cells across the integrated dataset were removed. The remaining gene-expression counts were library-size normalized (target sum = 10^4^) and log-transformed. Subsequently, the top 3500 highly variable genes were identified, and principal component analysis (PCA) was performed on the resulting variable-gene set, retaining the first 50 principal components for downstream analysis.

To account for the technical variation arising from study-specific effects and sequencing chemistry, cells were harmonised using the Harmony package in Python, [61] employing the composite study/chemistry batch annotation described in Section 5.2. Importantly, the harmonisation procedure did not modify the underlying gene-expression values; instead, it generated a batch-corrected low-dimensional representation that was subsequently used for neighbourhood graph construction, clustering, and the identification of shared cell populations across datasets. A nearest-neighbour graph (*n*_neighbours_ = 30) was constructed from the harmonised latent space, followed by Leiden clustering (resolution=10). Cell identities were subsequently assigned using the CellTypist package in Python, [45] which utilises reference transcriptomic signatures, in this case derived from healthy adult human skin, [46] to provide consistent cell-type annotation of the integrated clusters. The expression patterns of the eight consensus biomarkers were subsequently examined across the annotated cell populations and compared between cells originating from keloid and non-keloid samples.

To move beyond qualitative visualisation and quantitatively assess cell-type-specific differential expression, a pseudo-bulk differential gene expression analysis was performed for each annotated cell population to identify the cell populations most strongly contributing to the transcriptomic signatures associated with keloid pathology. For a given cell type, genes were retained if they accumulated at least 100 total raw counts across all samples and were detected in at least 35% of samples. Gene expression levels were then pseudo-bulked at the sample level by summing the raw counts across all cells belonging to the corresponding cell type within each sample. Differential gene expression analysis was subsequently performed on the resulting cell-type-specific pseudo-bulk profiles using DESeq2, yielding cell-type-specific estimates of *log*_2_-fold change and multiple-testing-adjusted (p)-values for all retained genes, from which the statistics for the identified biomarkers were extracted. As in Section 5.9, the analysis controlled for the composite study/chemistry batch effect while preserving the biological condition of interest (keloid versus non-keloid).

## 6 Acknowledgements

The authors would like to thank Paul Daniel, Kristin O’Malley, and Tim Hou (Qiagen) for their assistance and valuable discussions regarding the Ingenuity Pathway Analysis (IPA) performed in this study. The authors also thank Tracy Tabib (University of Pittsburgh) for helpful discussions regarding the identification of relevant skin scRNA-seq datasets and the interpretation of sequencing chemistry differences across studies. The computational analyses presented in this work were performed using the University of Dundee High Performance Computing (HPC) facility.

## Supplementary Material

**Table S1:**
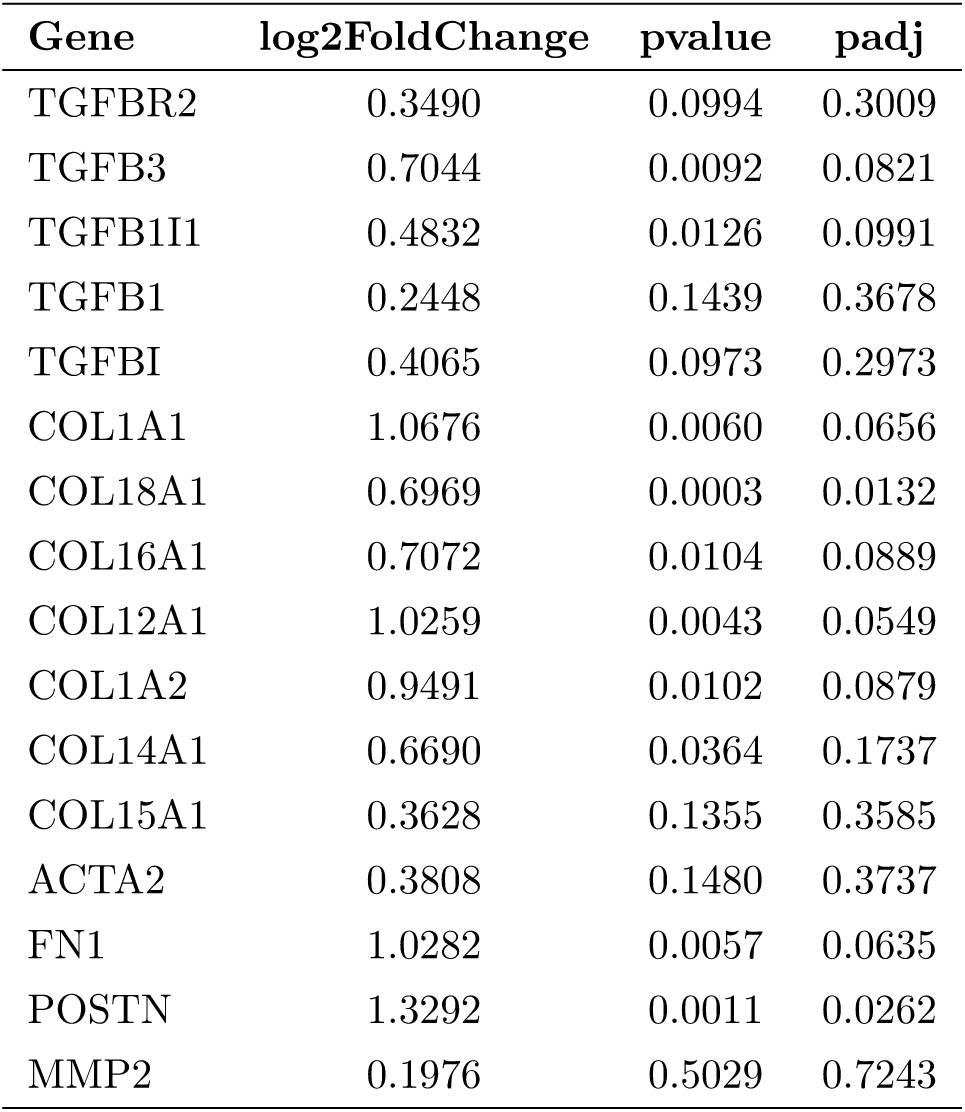
The Differential Gene Expression analysis results when using DESeq2 on the original 81 samples. for common target genes implicated in Keloid pathology.

| Gene | log2FoldChange | pvalue | padj |
| --- | --- | --- | --- |
| TGFBR2 | 0.3490 | 0.0994 | 0.3009 |
| TGFB3 | 0.7044 | 0.0092 | 0.0821 |
| TGFB1I1 | 0.4832 | 0.0126 | 0.0991 |
| TGFB1 | 0.2448 | 0.1439 | 0.3678 |
| TGFBI | 0.4065 | 0.0973 | 0.2973 |
| COL1A1 | 1.0676 | 0.0060 | 0.0656 |
| COL18A1 | 0.6969 | 0.0003 | 0.0132 |
| COL16A1 | 0.7072 | 0.0104 | 0.0889 |
| COL12A1 | 1.0259 | 0.0043 | 0.0549 |
| COL1A2 | 0.9491 | 0.0102 | 0.0879 |
| COL14A1 | 0.6690 | 0.0364 | 0.1737 |
| COL15A1 | 0.3628 | 0.1355 | 0.3585 |
| ACTA2 | 0.3808 | 0.1480 | 0.3737 |
| FN1 | 1.0282 | 0.0057 | 0.0635 |
| POSTN | 1.3292 | 0.0011 | 0.0262 |
| MMP2 | 0.1976 | 0.5029 | 0.7243 |

**Table S2:** Sensitivity and speciticity for keloid versus non-keloid discrimination across the ML frameworks. For multiclass classifiers, predictions were collapsed into keloid and non-keloid categories for evaluation purposes. Values are reported as mean ± standard deviation across outer folds.

| Algorithm | Sensitivity | Specificity |
| --- | --- | --- |
| Bin-LR | 0.89 ± 0.12 | 0.90 ± 0.08 |
| Bin-SVM | 0.91 ± 0.07 | 0.82 ± 0.14 |
| Multi-LR | 0.72 ± 0.20 | 0.95 ± 0.08 |
| Multi-SVM | 0.79 ± 0.16 | 0.89 ± 0.09 |
| Multi-RF | 0.82 ± 0.14 | 0.89 ± 0.12 |

**Figure S1:**
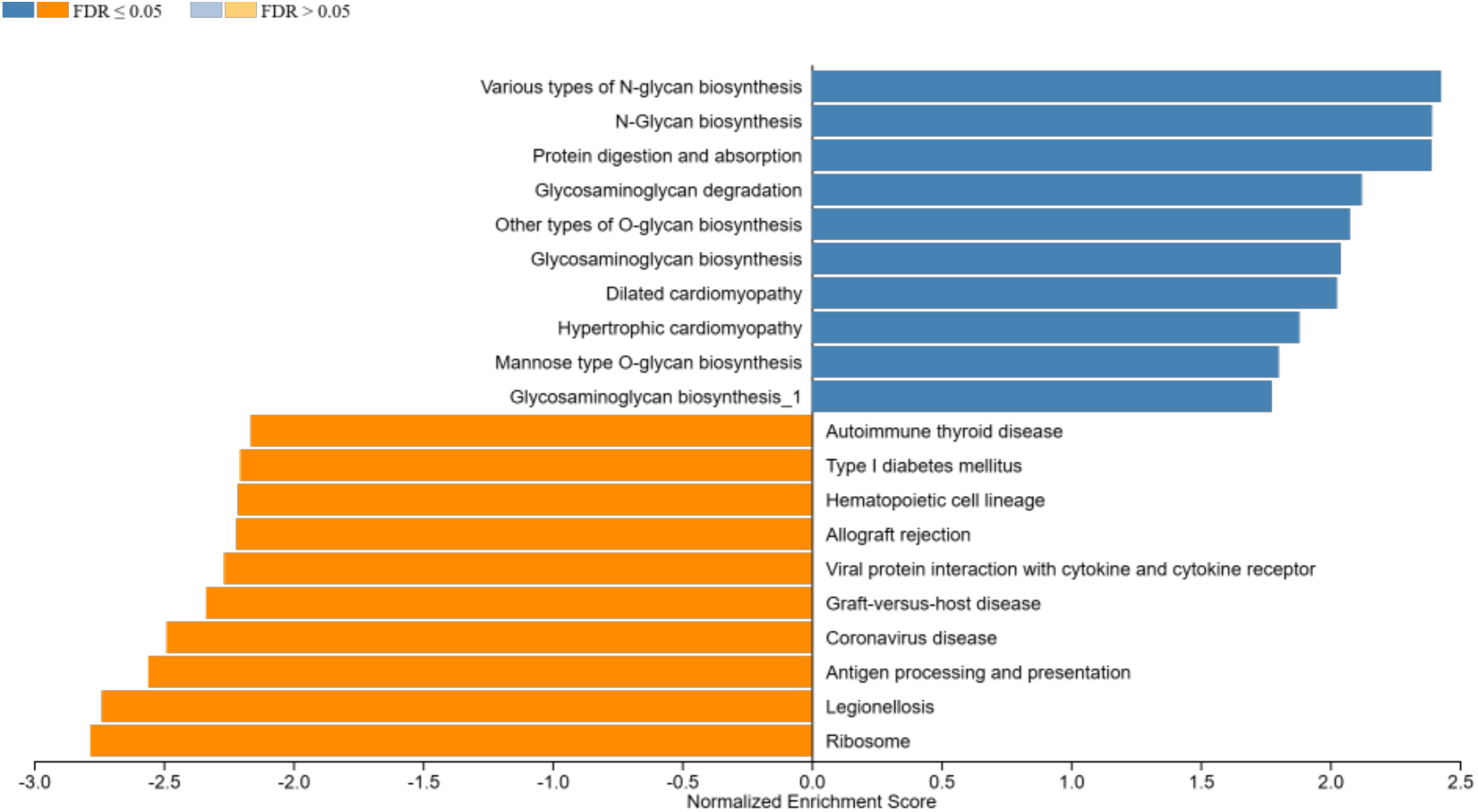
Gene Set Enrichment Analysis (GSEA) [21] of differentially expressed genes. Enriched pathways are ordered by Normalised Enrichment Score (NES). Dark range and Blue bars indicate statistically significant (*FDR* ≤ 0.05) enriched patwhays, respectively.

